# MorphoANOVA: Dissecting Histology–Gene Expression Relationships in Spatial Transcriptomics

**DOI:** 10.64898/2026.11.05.764699

**Authors:** Po-Wei Chen, Guanao Yan, Shan Yu, Jingyi Jessica Li

## Abstract

Spatial transcriptomics (ST) measures the expression of hundreds to thousands of genes within tissue architecture, often accompanied by histology images. However, it remains challenging to systematically characterize which genes exhibit associations with tissue morphology, whether these associations vary spatially, and which genes show additional spatial expression patterns beyond those captured by morphology. Here we introduce MorphoANOVA, a gene-wise statistical framework for characterizing morphology– expression relationships across genes measured by ST. Using nested spatially varying coefficient models and sequential ANOVA, MorphoANOVA assesses slide-wide associations with an H&E-derived morphology feature, location-specific changes in these associations, and additional spatial variation after accounting for the selected feature. The resulting statistics enable gene prioritization and pathway enrichment analysis, while spatial component maps support the interpretation of individual genes. Simulations demonstrate MorphoANOVA’s ability to distinguish the modeled components and recover their spatial patterns. Applications to human cortex and breast cancer identify biological programs associated with different morphology–expression patterns and reveal distinct relationships with cell composition. Across three HER2-positive breast cancer patients, invasive-cancer marker genes consistently exhibit stronger evidence of location-specific morphology associations. MorphoANOVA thus provides a framework for systematically interpreting spatial gene-expression patterns in relation to tissue morphology.

## Introduction

Spatial transcriptomics (ST) technologies enable the measurement of hundreds to tens of thousands of genes while preserving their spatial locations within tissue architecture, providing insights into spatial cellular organization and function. Advances in spatial resolution have expanded ST from multi-cell capture spots, as in early spatial barcoding approaches [1], to single-cell resolution with methods such as Slide-seq [2] and HDST [3], and even to subcellular levels, as achieved by high-resolution in situ hybridization methods like MERFISH [4], seqFISH [5], and Xenium [6]. Hematoxylin and eosin (H&E)-stained histology images, routinely generated in clinical and research workflows, provide complementary structural and morphological information and have also been shown to be associated with spatial gene expression patterns [7]. These observations suggest that tissue morphology may offer important clues to spatial molecular organization.

Morphology–expression relationships may differ across genes and tissue regions. Even within the same cell type, transcriptional programs can vary across tissue regions because of differences in microenvironmental signals and cellular states [8], as well as tissue-specific regulation [9, 10]. Consequently, histologically similar regions may exhibit distinct expression patterns, while the association between a morphology feature and a gene’s expression may be strong in some regions but weak or absent in others. These observations motivate a systematic approach to determining, for each gene, whether its expression is associated with tissue morphology, whether this association varies spatially, and whether additional spatial expression patterns remain after accounting for the selected morphology feature.

Numerous computational methods have been developed to jointly analyze ST and histological images, broadly focusing on expression prediction or multimodal integration [11]. Prediction-based approaches aim to predict gene expression from histology, either in unmeasured regions or at higher spatial resolution. Early methods such as ST-Net [12] use convolutional neural networks, while Hist2ST [13] combines image features with spatial context for expression prediction. Subsequent studies include BLEEP [14], which uses contrastive bimodal embeddings for prediction, and iSTAR [15], which uses a self-supervised vision Transformer to extract hierarchical histology features and predict gene expression at higher spatial resolution. Integration-based approaches combine ST and histology, often to improve spatial domain identification. SpaCell [16] and stLearn [17] both use pretrained ResNet50 [18] to extract histology features. SpaCell embeds these jointly with gene expression, whereas stLearn combines them with spatial neighborhood information to adjust expression values. SpaGCN [19] integrates ST and histology through graph convolutional networks to identify spatial domains. Other integration methods, including conST [20] and ConGI [21], use contrastive learning to integrate ST and histology for spatial clustering.

Despite these advances, existing methods provide limited tools for systematically characterizing how morphology– expression relationships vary across tissue and across genes. Early studies, such as those by Badea et al. [22] and ImageCCA [7], associated histological image features with gene expression but operated mainly at the whole-slide or tissue-sample level. More recent methods, such as SPADE [23] and MorphLink [24], identify morphology-associated genes, with MorphLink additionally quantifying local morphology–expression concordance within tissue subregions. However, these methods do not provide a unified gene-wise framework for separately assessing slide-wide morphology associations, location-specific changes in these associations, and additional spatial expression patterns after accounting for the selected morphology feature. Such a framework would enable researchers to compare morphology–expression relationships across genes and identify where distinct spatial patterns occur within tissue.

Here we introduce MorphoANOVA, a statistical framework for systematically characterizing morphology–expression relationships across genes measured by spatial transcriptomics. For each gene, MorphoANOVA uses sequential analysis of variance (ANOVA) of nested spatially varying coefficient models (SVCMs) to assess three questions: whether gene expression is associated with a selected H&E-derived morphology feature across the tissue, whether this association varies by location, and whether additional spatial expression patterns remain after accounting for the selected feature. MorphoANOVA assesses the statistical evidence for these patterns, summarizes the corresponding sequential improvements in adjusted *R*^2^, and maps their spatial distributions. These results support two complementary applications: prioritizing genes and biological pathways with distinct morphology–expression relationships, and identifying tissue regions where morphology-associated and additional spatial expression patterns occur. Applications to human cortex and breast cancer demonstrate how these patterns can relate differently to cell composition and tumor states.

## Results

### Method overview

MorphoANOVA systematically characterizes morphology–expression relationships across genes measured by spatial transcriptomics. As shown in Fig. 1a, the framework takes paired H&E histology and spatial gene-expression measurements as input and extracts a morphology feature at each spatial location. For each gene, MorphoANOVA fits four SVCMs (Fig. 1b). Starting from an intercept-only model, the sequence adds a slide-wide H&E effect, a location-specific H&E effect capturing spatial changes in the association, and additional spatial variation after accounting for the selected H&E feature. The slide-wide and location-specific H&E effects together define the total H&E effect. Sequential ANOVA assesses statistical evidence for the three patterns: *F*_21_ tests the slide-wide H&E effect, *F*_32_ tests the additional location-specific H&E effect, and *F*_43_ tests spatial variation beyond H&E. Sequential changes in adjusted *R*^2^ summarize the corresponding improvements in model fit. Further methodological details are provided in Methods.

**Figure 1:**
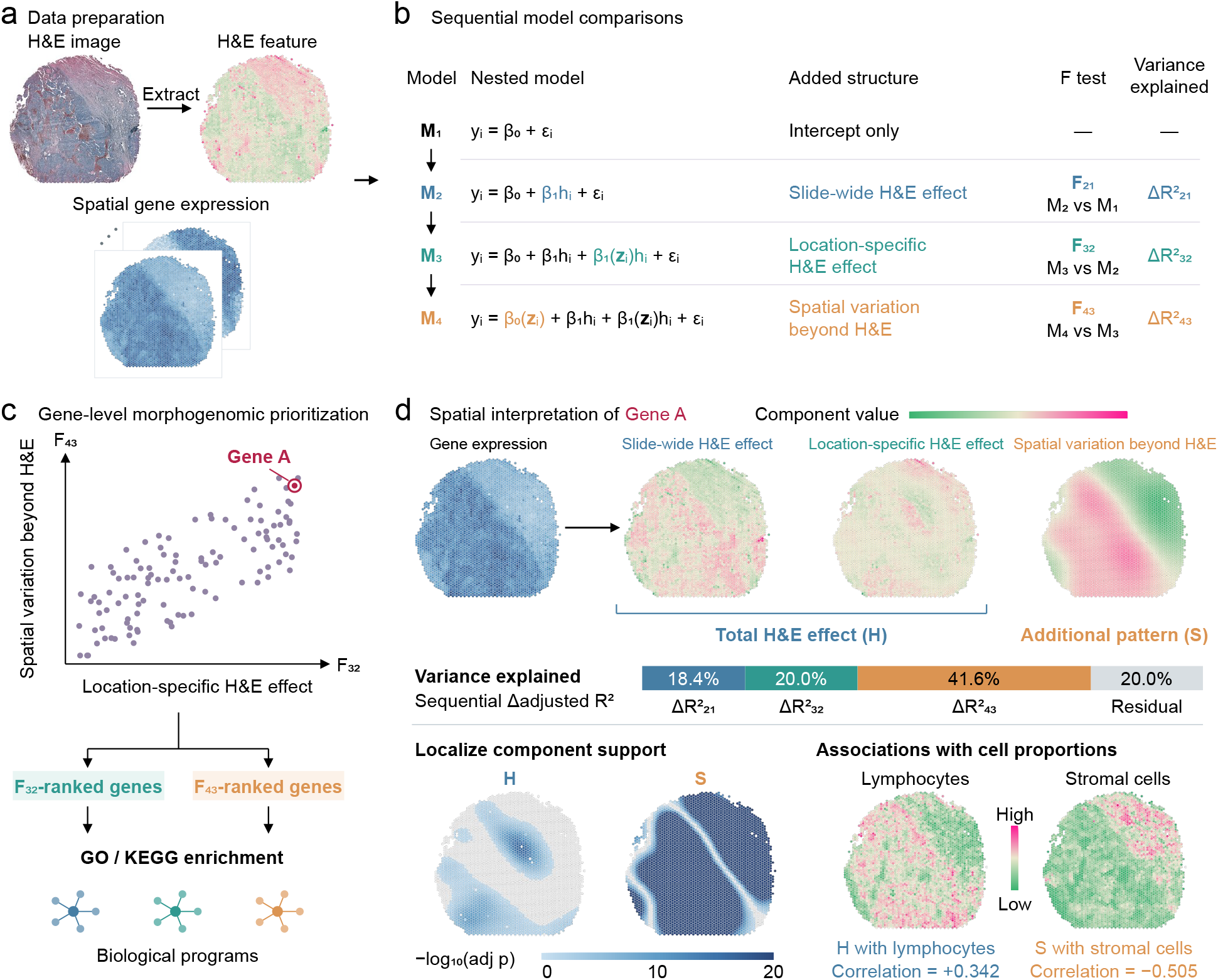
Workflow of MorphoANOVA. (**a**) Extraction of an H&E-derived morphology feature paired with spatial gene-expression measurements. (**b**) Four nested models assess slide-wide H&E effects (*F*_21_), location-specific H&E effects (*F*_32_), and spatial variation beyond the selected H&E feature (*F*_43_). Sequential changes in adjusted *R*^2^ summarize the corresponding improvements in model fit. (**c**) Gene prioritization by *F*_32_- and *F*_43_-based rankings, followed by GO and KEGG enrichment analyses to identify associated biological programs. (**d**) Spatial interpretation of an individual gene using fitted component maps. Adjusted *p*-value maps identify regions with statistical support for the total H&E effect or spatial variation beyond H&E, while correlations with cell-type proportions characterize their cellular associations.

MorphoANOVA supports two complementary applications (Fig. 1c,d). First, at the gene level, it uses *F*_32_ and *F*_43_ to characterize morphology–expression relationships and prioritize genes with strong location-specific H&E effects or additional spatial variation. These genes are analyzed for enrichment of Gene Ontology (GO) Biological Process terms and Kyoto Encyclopedia of Genes and Genomes (KEGG) pathways to identify associated biological programs. Second, for individual genes, MorphoANOVA maps the fitted slide-wide H&E effect, location-specific H&E effect, and spatial variation beyond H&E, and identifies tissue regions where the total H&E effect or additional spatial variation is statistically supported. By correlating the fitted spatial components with cell-type proportions, MorphoANOVA identifies which cell types are associated with the H&E-related and additional spatial expression patterns.

### MorphoANOVA distinguishes and recovers spatial expression patterns in simulations

We used simulations with known spatial expression patterns to evaluate MorphoANOVA’s ability to distinguish and recover the three modeled components: the slide-wide H&E effect, location-specific H&E effect, and spatial variation beyond H&E. To preserve realistic spatial geometry and morphology patterns, we simulated gene expression at 3,455 observed spots from an ovarian cancer Visium dataset, using its H&E image to derive the morphology covariate. This dataset was also analyzed in the Supplementary Information. Gene expression for four gene classes was generated from nested models, starting with an intercept-only model and sequentially adding slide-wide H&E, location-specific H&E, and spatial variation beyond H&E (Fig. 2a). Each setting was evaluated over 500 replicates.

**Figure 2:**
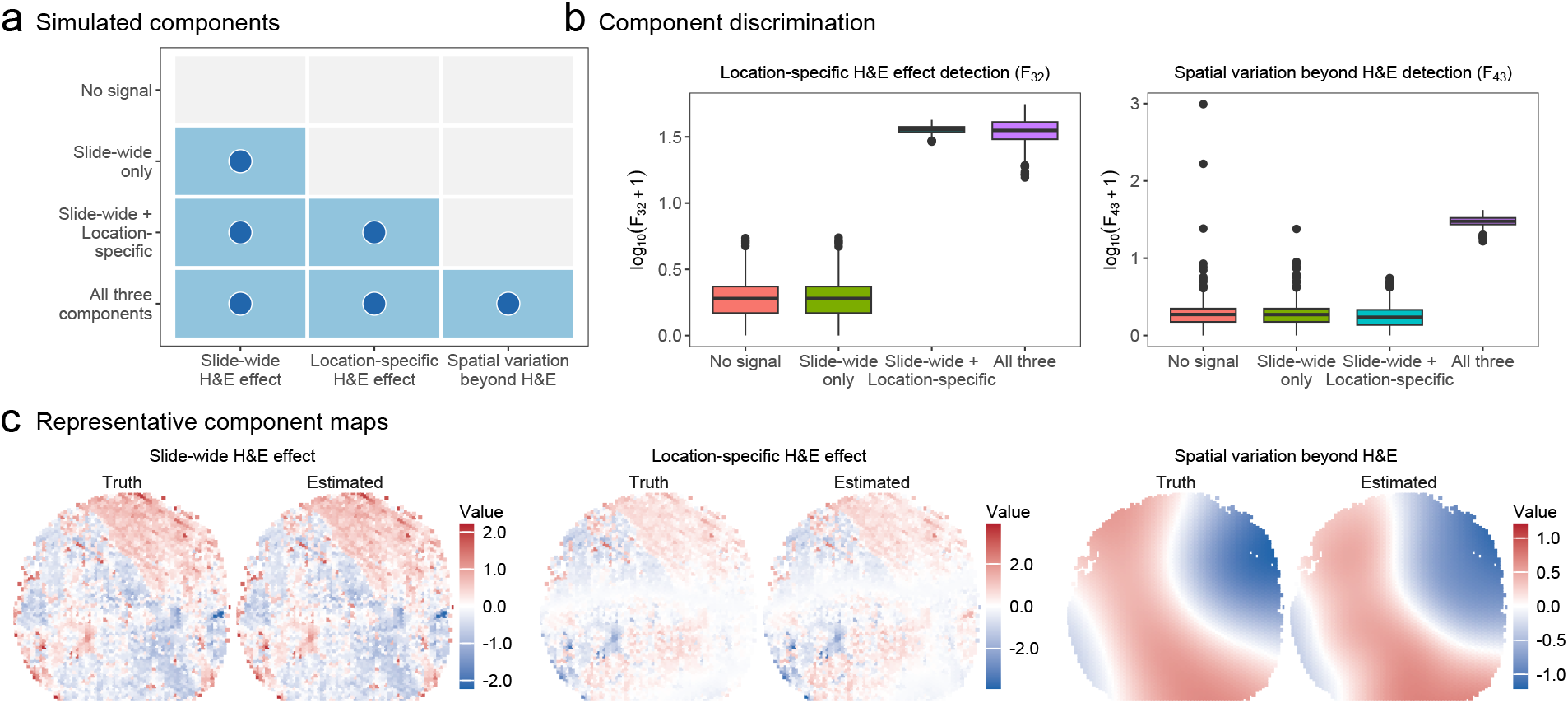
MorphoANOVA distinguishes and recovers simulated spatial expression patterns. (**a**) Four simulated gene classes based on nested models: no modeled signal; slide-wide H&E effect only; slide-wide and location-specific H&E effects; and all three components. (**b**) Distributions of *F*_32_ and *F*_43_ across the four gene classes over 500 replicates. (**c**) True and estimated spatial patterns of the three components. Panels (**b**,**c**) use *n* = 3455, with an SNR of 0.25 for each active component.

MorphoANOVA distinguished the simulated gene classes and recovered the underlying spatial patterns. Specifically, *F*_32_ increased in the presence of a location-specific H&E effect, whereas *F*_43_ increased only when additional spatial variation beyond H&E was present (Fig. 2b). The estimated components also reproduced the major spatial patterns of their simulated counterparts (Fig. 2c). The displayed replicate was selected as the median among 500 replicates based on the Pearson correlation between true and estimated spatial variation beyond H&E.

Additional simulations assessed statistical performance across sample sizes of 500–3455 spatial locations, signal-to-noise ratios (SNRs) of 0.01–0.25 for the three components, and correlations of 0–0.95 between the H&E feature and spatial variation beyond H&E. Type-I error rates were generally close to the nominal 5% level, with 100% detection power in most tested settings. Sequential improvements in adjusted *R*^2^ were generally consistent with the corresponding signal-based reference values (Supplementary Fig. 1a–c). Additional simulation details are provided in Methods.

### Linking cortical transcriptional programs to H&E effects and spatial variation beyond H&E

We first applied MorphoANOVA to the human cortex to examine how its laminar organization is reflected in morphology– expression relationships. We analyzed human dorsolateral prefrontal cortex (DLPFC) sample 151673, comprising measurements of 33,538 genes across 3,639 Visium spots [25]. After filtering for expression variability, 8,919 genes were retained. At each spatial location *i*, we define a scalar H&E-derived morphology covariate *h*_*i*_^raw^, whose construction is specified by the user according to the dataset and scientific question. In this study, we used an H&E intensity feature extracted from image patches centered on each spot to summarize local staining appearance, one aspect of the visual information considered in histological assessment (Fig. 3a). The six cortical layers and white matter provided a structured setting for examining which spatial expression patterns were associated with the selected H&E feature and which remained after accounting for it.

**Figure 3:**
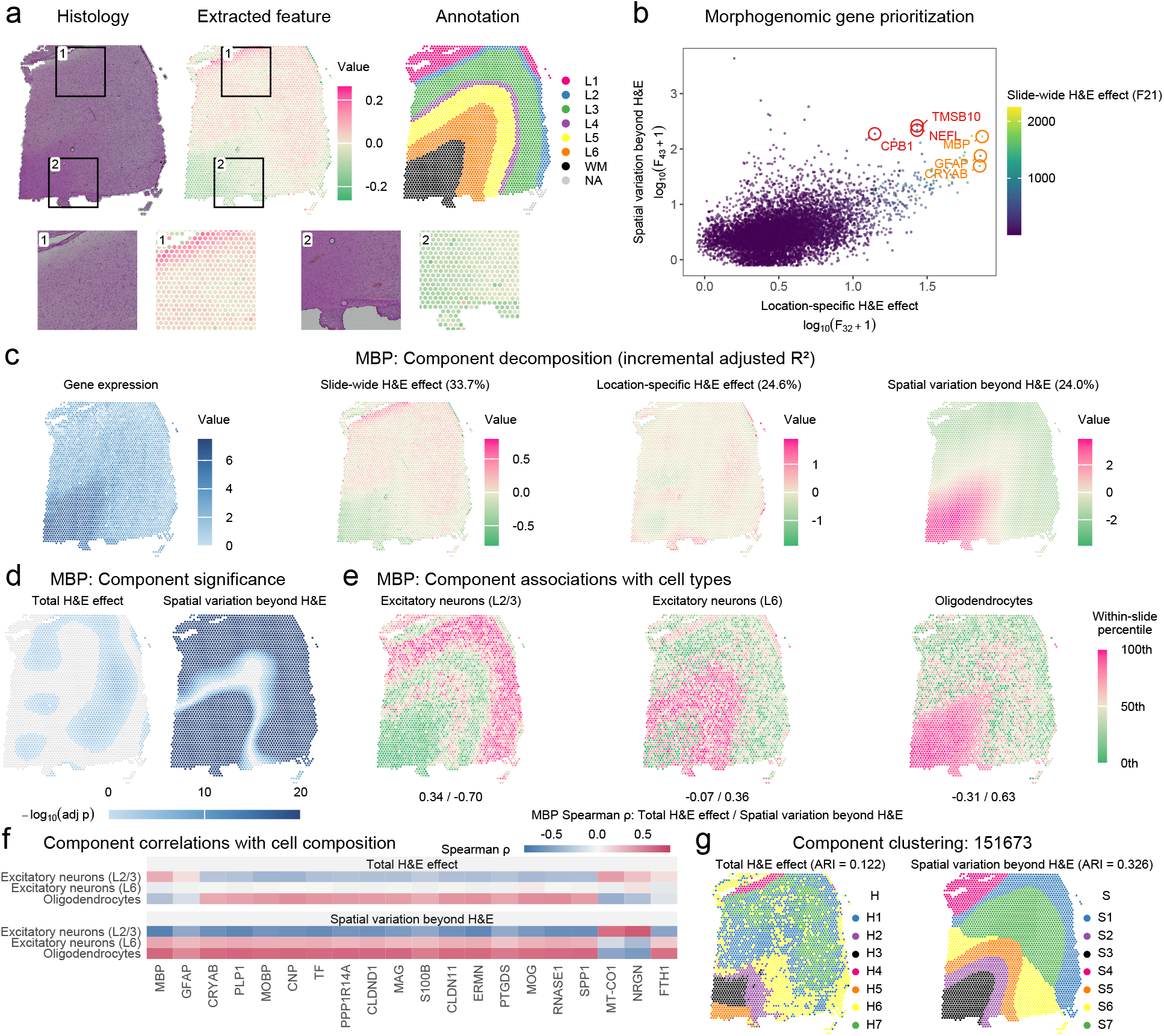
MorphoANOVA characterizes morphogenomic patterns in human cortex tissue. (**a**) H&E image, extracted morphology feature, and cortical-layer annotations for DLPFC sample 151673, with enlarged views of two marked regions. (**b**) Morphogenomic gene prioritization of log_10_(*F*_32_ + 1) (horizontal axis) and log_10_(*F*_43_ + 1) (vertical axis), colored by *F*_21_. Orange and red labels mark the top three genes in the *F*_32_- and *F*_43_-based rankings, respectively. (**c**) Observed *MBP* expression and three fitted components; percentages indicate sequential increases in adjusted *R*^2^. (**d**) BH-adjusted pointwise *p*-value maps for the total H&E effect and spatial variation beyond H&E. Spots with adjusted *p >* 0.05 are gray. (**e**) Estimated cell-type proportions shown as within-slide percentiles. Paired values give Spearman correlations with the *MBP* total H&E effect and spatial variation beyond H&E, respectively. (**f**) Corresponding correlations for 20 of the 102 genes shared between the two top-200 lists. (**g**) Spatial clustering of spots using the total H&E effect and spatial variation beyond H&E across the *F*_32_- and *F*_43_-based top-200 gene lists, respectively.

We first prioritized genes with strong location-specific H&E effects or spatial variation beyond H&E. For each gene, we summarized the corresponding model-comparison statistics *F*_32_ and *F*_43_ (Fig. 3b). After requiring a sequential increase in adjusted *R*^2^ of at least 0.03 for each component, we ranked genes by *F*_32_ and *F*_43_, respectively, and selected the top 200 genes from each ranking for functional enrichment analysis (top six shown in Supplementary Fig. 2a). GO enrichment of genes with strong location-specific H&E effects highlighted axon ensheathment, oligodendrocyte differentiation, and substantia nigra development. In contrast, genes with strong spatial variation beyond H&E were enriched for proton transmembrane transport, intermediate filament organization, and oxidative phosphorylation. KEGG enrichment identified oxidative phosphorylation and Parkinson disease in both gene lists (Supplementary Fig. 2b).

We next examined *MBP*, which ranked highly in both gene lists, to illustrate how MorphoANOVA characterizes an individual gene’s spatial expression pattern. The full model achieved an adjusted *R*^2^ of 82.4%, with sequential increases of 33.7, 24.6, and 24.0 percentage points upon adding the slide-wide H&E effect, location-specific H&E effect, and spatial variation beyond H&E, respectively. The location-specific H&E effect showed a band-like pattern consistent with cortical lamination, whereas spatial variation beyond H&E extended across much of the tissue section (Fig. 3c). Significance maps identified regions with statistical support for the total H&E effect and spatial variation beyond H&E (Fig. 3d). To characterize their cellular associations, we calculated Spearman correlations between the fitted spatial components and estimated cell-type proportions across spots (Fig. 3e). The total H&E effect correlated with layer 2/3 excitatory-neuron proportions (*ρ* = 0.34) but showed little correlation with layer 6 excitatory-neuron proportions (*ρ* = −0.07). In contrast, spatial variation beyond H&E showed stronger correlations in absolute value with both cell types (*ρ* = −0.70 and 0.36, respectively). Consistent with the established association between *MBP* and oligodendrocytes [26], spatial variation beyond H&E was also more strongly correlated with oligodendrocyte proportions (*ρ* = 0.63) than the total H&E effect (*ρ* = −0.31).

We further examined correlations between the fitted spatial components and estimated cell-type proportions for the 102 genes shared by the two top-200 lists (Fig. 3f; remaining genes in Supplementary Fig. 3a). For layer 6 excitatory neurons, correlations with the total H&E effect were consistently weak, whereas spatial variation beyond H&E showed stronger correlations for many genes. For layer 2/3 excitatory neurons and oligodendrocytes, both components were correlated with their estimated proportions, but the absolute correlations were generally stronger for spatial variation beyond H&E. Together, these findings indicate that the selected H&E feature captures only part of the spatial transcriptional organization of the cortex, with additional spatial expression patterns showing stronger associations with certain cortical cell populations.

We next clustered spots based on the fitted total H&E effect and spatial variation beyond H&E for the *F*_32_- and *F*_43_-based top-200 gene sets, respectively, to examine heterogeneity within anatomical compartments (cortical layers L1–L6 and white matter) and cortical layer organization (Fig. 3a,g). Within annotated white matter, spots assigned to H2, H3, and H5 had mean estimated oligodendrocyte proportions of 52.0%, 62.6%, and 67.5%, respectively. H4 was located within L1 and had a higher mean estimated astrocyte proportion than the remaining L1 spots (23.1% versus 10.6%). Supporting this interpretation, three additional astrocyte markers showed higher mean observed expression in H4 than in the remaining L1 spots (Supplementary Fig. 3b). In contrast, spatial variation beyond H&E showed greater agreement with anatomical annotations than the total H&E effect (adjusted Rand index (ARI), 0.326 versus 0.122). However, S7 spanned L3–L6 rather than a single cortical layer. The fitted spatial patterns of *SCGB2A2* and *SCGB1D2* suggested that they contributed to the S6–S7 separation. Removing these genes increased the ARI to 0.380 (Supplementary Fig. 3c), consistent with a previous STModule study noting that they are not layer-specific markers [27]. These results show that morphology-associated patterns delineated subregions within anatomical compartments, whereas spatial variation beyond H&E more closely reflected cortical laminar organization.

### Distinguishing morphogenomic components of gene expression at cellular resolution

We next applied MorphoANOVA to a Xenium breast cancer dataset to examine morphology–expression relationships at single-cell resolution. The dataset comprises measurements of 313 genes across 167,780 segmented cells, together with a high-resolution H&E image [6]. To obtain a data-driven representation of tissue morphology, we extracted hierarchical histology features using iSTAR [15] and used their first principal component as the morphology covariate (Fig. 4a). We then examined which genes were associated with this learned feature, how these associations varied spatially, and which additional spatial expression patterns remained after accounting for it.

**Figure 4:**
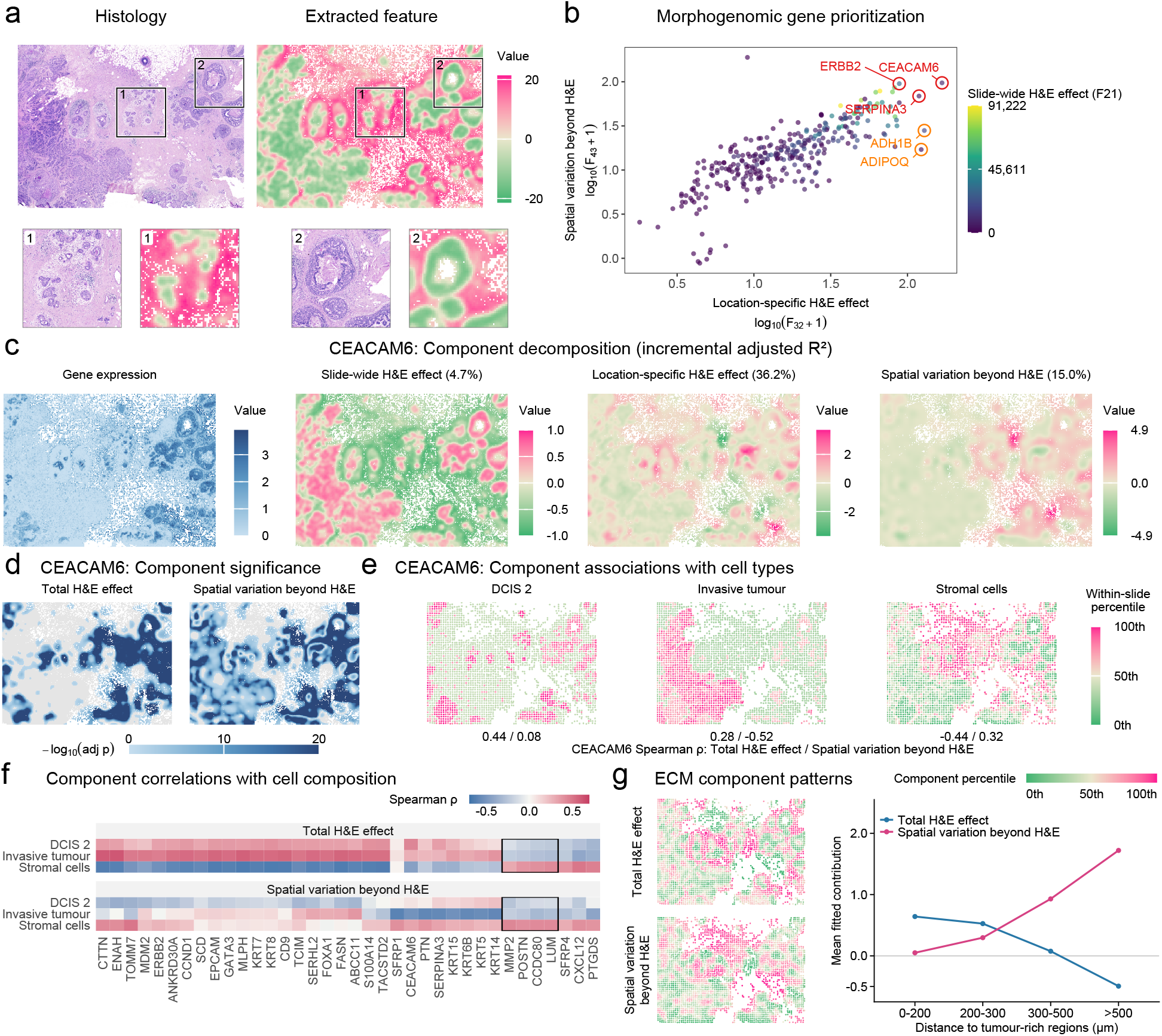
MorphoANOVA characterizes morphology–expression relationships in breast cancer at single-cell resolution. (**a**) H&E image and extracted morphology feature, with enlarged views of two marked regions. (**b**) Morphogenomic gene prioritization using log_10_(*F*_32_ + 1) (horizontal axis) and log_10_(*F*_43_ + 1) (vertical axis), colored by *F*_21_. Orange and red labels mark the top three genes in the *F*_32_- and *F*_43_-based rankings, respectively. (**c**) Observed *CEACAM6* expression and three fitted components; percentages indicate sequential increases in adjusted *R*^2^. (**d**) BH-adjusted pointwise *p*-value maps for the total H&E effect and spatial variation beyond H&E gray indicates adjusted *p >* 0.05. (**e**) DCIS 2, invasive-tumour, and stromal-cell proportions shown as within-slide percentiles. Paired values indicate Spearman correlations with the *CEACAM6* total H&E effect and spatial variation beyond H&E, respectively. Corresponding correlations for the 36 genes shared by the two top-50 lists. Outlined columns indicate the four ECM organization genes examined in (**g**). (**g**) Spatial patterns of the total H&E effect and spatial variation beyond H&E, averaged across four ECM organization genes (*CCDC80, LUM, MMP2*, and *POSTN*). The distance plot shows mean fitted contributions in stromal-rich bins at increasing distances from tumour-cell-rich regions. Analyses in (**e**–**g**) used spatial bins of 100 × 100 *µ*m; correlations in (**e**,**f**) were calculated across 3,267 bins.

We first prioritized genes with strong location-specific H&E effects or spatial variation beyond H&E. For each gene, we summarized the corresponding model-comparison statistics *F*_32_ and *F*_43_ (Fig. 4b). After requiring a sequential increase in adjusted *R*^2^ of at least 0.03 for each component, we ranked genes by *F*_32_ and *F*_43_, respectively, and selected the top 50 from each ranking for functional enrichment analysis (top six shown in Supplementary Fig. 4a). GO enrichment identified supramolecular fiber organization and neuron projection development in both gene lists, with tissue development and axon development among the leading terms in the *F*_32_- and *F*_43_-based lists, respectively. KEGG enrichment identified Cadherin signaling and Cornified envelope formation in both lists, with Proteoglycans in cancer additionally enriched among genes with strong location-specific H&E effects (Supplementary Fig. 4b).

We next examined *CEACAM6*, which ranked first in both gene lists, to illustrate how MorphoANOVA characterizes an individual gene’s spatial expression pattern. The full model achieved an adjusted *R*^2^ of 55.9%, with sequential increases of 4.7, 36.2, and 15.0 percentage points upon adding the slide-wide H&E effect, location-specific H&E effect, and spatial variation beyond H&E, respectively. The fitted components showed distinct spatial patterns (Fig. 4c), with regions of statistical support shown in Fig. 4d. To examine their relationships with tissue composition, we calculated Spearman correlations between the fitted spatial components and estimated cell-type proportions across spatial bins (Fig. 4e). The total H&E effect was more strongly correlated with the proportion of cells annotated as ductal carcinoma in situ (DCIS) category 2 (*ρ* = 0.44) than spatial variation beyond H&E (*ρ* = 0.08). Conversely, spatial variation beyond H&E showed a stronger absolute correlation with invasive-tumour proportions (*ρ* = −0.52) than the total H&E effect (*ρ* = 0.28). The two components were also correlated with stromal-cell proportions (*ρ* = −0.44 and 0.32, respectively). Thus, the total H&E effect for *CEACAM6* was more closely associated with DCIS 2, whereas spatial variation beyond H&E showed a stronger association with invasive-tumour composition.

We further examined correlations between the fitted spatial components and cell-category proportions for the 36 genes shared by the two top-50 lists (Fig. 4f). Across these genes, the total H&E effect generally showed stronger absolute correlations with DCIS 2, invasive-tumour, and stromal-cell proportions than spatial variation beyond H&E.

However, spatial variation beyond H&E showed stronger absolute correlations with invasive-tumour proportions for some genes, including *CEACAM6*. Results for 17 additional cell populations are shown in Supplementary Fig. 5. These findings show that the learned H&E feature captures spatial expression patterns associated with tumor and stromal composition, while additional spatial patterns beyond this feature provide complementary information for certain genes.

**Figure 5:**
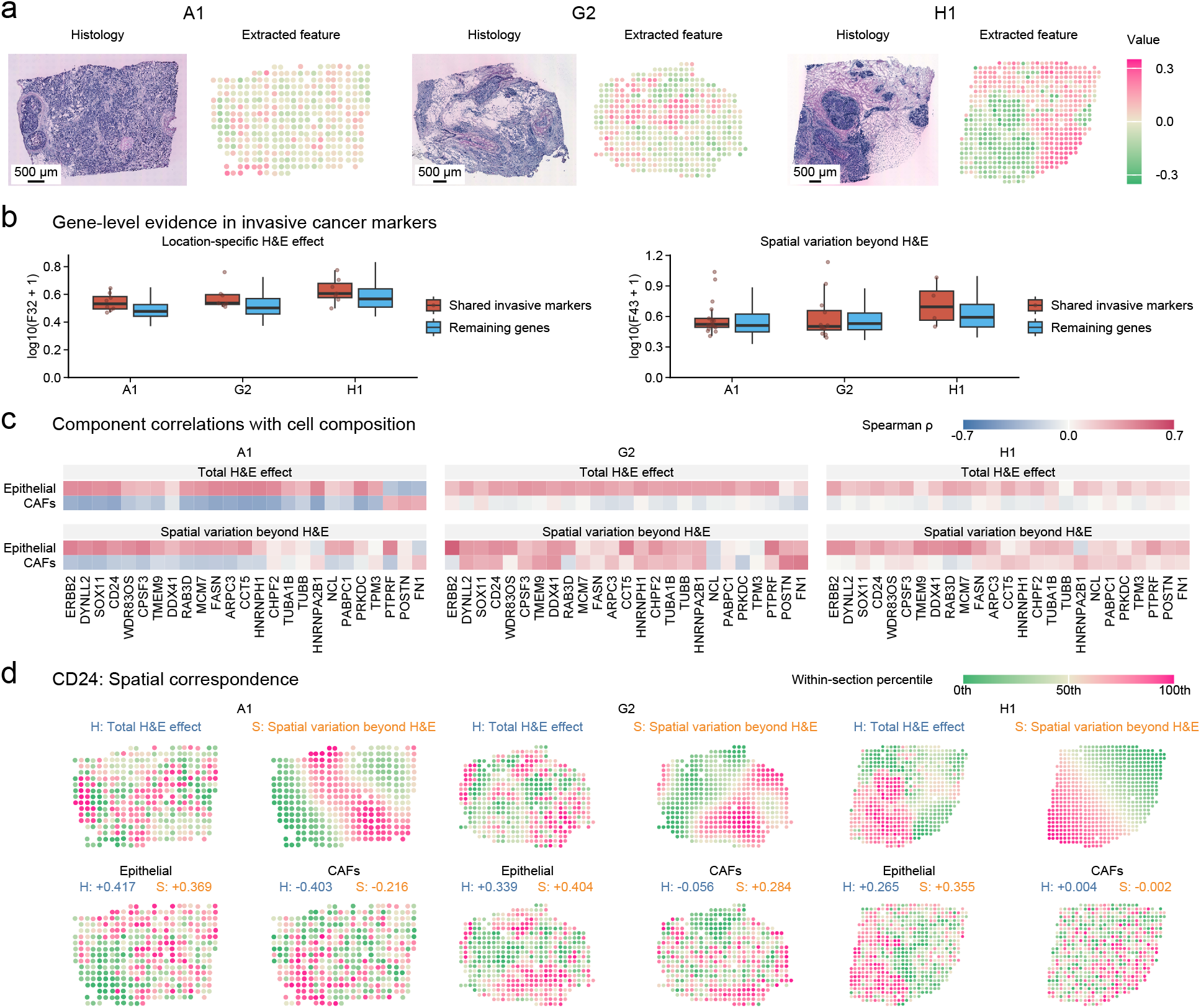
Shared and patient-specific morphology–expression patterns in HER2^+^ breast cancer. (**a**) H&E images and extracted morphology features for sections A1, G2, and H1. (**b**) Distributions of *F*_32_ and *F*_43_ for the 25 shared invasive-cancer marker genes and the remaining genes in each section. (**c**) Spearman correlations between the fitted total H&E effect or spatial variation beyond H&E and estimated epithelial-cell and CAF proportions across spots, shown for the 25 shared markers. (**d**) Spatial maps of the two components for *CD24*, alongside estimated epithelial-cell and CAF proportions. Annotations indicate their Spearman correlations within each section.

To characterize the spatial relationship between stromal-associated genes and the selected H&E feature, we focused on four genes involved in extracellular matrix (ECM) organization (outlined in Fig. 4f). Within each spatial bin, we averaged the fitted total H&E effect and spatial variation beyond H&E across the four genes. Both components were positively correlated with stromal-cell proportions (Spearman *ρ* = 0.393 and 0.431, respectively) but showed distinct spatial patterns (Fig. 4g). In stromal-rich bins, the H&E-associated contribution was higher near the tumour, whereas spatial variation beyond H&E was higher in more distant stroma. These results show how MorphoANOVA distinguishes morphology-guided patterns from additional spatial variation beyond morphology.

### MorphoANOVA reveals shared and patient-specific morphology–expression patterns in breast cancer

We next applied MorphoANOVA to HER2^+^ breast cancer data from the HER2ST study [28] to examine whether morphology–expression relationships were consistent across patients. We analyzed sections A1, G2, and H1 from three patients. These sections cover all tumor subtypes in the original cohort and were previously analyzed by MorphLink [24]. We used a mean-centered H&E intensity feature extracted from approximately 100 *µ*m-wide image patches at each spot as the morphology covariate (Fig. 5a). Through differential expression analysis, we identified 25 invasive-cancer marker genes shared across the three sections and compared their morphology–expression relationships.

The shared invasive-cancer markers had higher median *F*_32_ values than the remaining genes in all three sections, indicating stronger evidence for location-specific H&E effects, whereas differences in *F*_43_ were less consistent (Fig. 5b). We further examined correlations between the fitted spatial components and estimated cell-type proportions across spots. Both the total H&E effect and spatial variation beyond H&E were predominantly positively correlated with epithelial-cell proportions in all three sections. In contrast, correlations with cancer-associated fibroblast (CAF) proportions varied across sections. They were predominantly negative for both components in A1 and generally weak in H1. In G2, correlations with CAF proportions were weak for the total H&E effect but positive for spatial variation beyond H&E (Fig. 5c). Results for six additional cell types are shown in Supplementary Fig. 6.

We selected *CD24* to illustrate these patterns (Fig. 5d). Both components were positively correlated with epithelial-cell proportions in all three sections, but their relationships with CAF proportions differed. In G2, spatial variation beyond H&E was positively correlated with CAF proportions (*ρ* = 0.284), whereas the total H&E effect showed little correlation (*ρ* = −0.056). Both components were negatively correlated with CAF proportions in A1, with negligible correlations in H1. These results show that shared invasive-cancer markers exhibit consistently stronger evidence for location-specific H&E effects across patients, while their fitted spatial components show patient-specific relationships with tissue cell composition.

## Discussion

We developed MorphoANOVA, a framework for characterizing morphology–expression relationships in ST. Using nested SVCMs and sequential ANOVA, it assesses slide-wide H&E effects, location-specific changes, and additional spatial variation beyond morphology. Simulations showed accurate recovery of these patterns with near-nominal type-I error rates. Applications to human cortex and breast cancer revealed distinct biological programs and cell-type associations, while shared invasive-cancer markers showed stronger location-specific H&E effects but patient-specific relationships with cell composition.

MorphoANOVA complements existing approaches for jointly analyzing histology and spatial transcriptomics. Whereas prediction-based methods estimate gene expression from histology and integration-based methods combine the two modalities for tasks such as spatial clustering, MorphoANOVA focuses on systematically characterizing morphology–expression relationships across genes. Unlike morphology-linking approaches such as MorphLink, which quantifies local morphology–expression concordance within tissue subregions, MorphoANOVA jointly assesses slide-wide associations, location-specific changes in these associations, and additional spatial expression patterns after accounting for morphology. These analyses help researchers prioritize genes with distinct morphology–expression relationships and identify where these patterns occur within tissue.

MorphoANOVA can also be applied to different types of H&E-derived morphology features. We demonstrated this flexibility using image-intensity features in human cortex and HER2^+^ breast cancer, and a learned iSTAR-derived histology feature in Xenium breast cancer. In particular, MorphoANOVA provides a way to interpret learned histology features by identifying genes and tissue regions associated with these features, as well as additional spatial expression patterns that they do not capture. This capability complements existing image-feature extraction methods by providing gene-level and spatial interpretation of their relationships with gene expression.

There are several limitations to our study. First, MorphoANOVA analyzes one morphology feature at a time, whereas tissue morphology is multidimensional. Consequently, spatial variation beyond H&E is defined relative to the selected feature and may contain variation associated with other histological characteristics. Extending the framework to multiple morphology features would require additional modeling and regularization. Second, the sequential improvements in adjusted *R*^2^ depend on the specified order of model inclusion and should not be interpreted as unique contributions from independent biological sources. Finally, strong overlap between spatial terms (concurvity) may make location-specific H&E effects and spatial variation beyond H&E difficult to distinguish [29].

## Methods

### Input data and morphology covariates

MorphoANOVA takes as input spatial transcriptomics measurements, spatial coordinates, and an aligned H&E histology image from the same tissue section. For a section with *n* spatial locations and *G* genes, let *Y* ∈ R^*n×G*^ denote the gene-expression matrix and **z**_*i*_ ∈ Ω the two-dimensional coordinate of location *i*, where Ω is the tissue’s spatial domain. For each gene, let *y*_*i*_ denote its log-transformed expression at location *i*.

Let 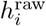 denote a scalar H&E-derived morphology feature at location *i*, obtained from image-intensity summaries, learned histology embeddings, or other image-analysis methods [15, 23, 24]. We center this feature within each section as 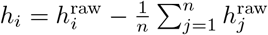.

### Nested SVCM specification and fitting

For each gene, MorphoANOVA fits four nested spatially varying coefficient models (SVCMs):

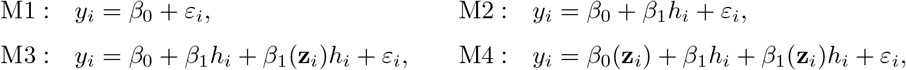

Where 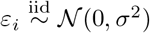. Starting from the intercept-only model M1, M2 i ntroduces a slide-wide H&E effect through *β*_1_, and M3 adds a location-specific H&E effect through *β*_1_(**z**_*i*_), with 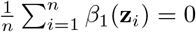. M4 replaces the constant intercept with a spatially varying intercept *β*_0_(**z**_*i*_), capturing additional spatial variation after accounting for the selected H&E feature. This sequence allows us to assess slide-wide morphology associations, location-specific changes in these associations, and spatial expression patterns remaining beyond the selected morphology feature. We fitted Gaussian generalized additive models with an identity link, representing spatially varying terms using penalized thin-plate regression splines and estimating smoothing parameters by restricted maximum likelihood (REML).

### Sequential ANOVA and incremental adjusted *R*^2^

We used sequential analysis of variance (ANOVA) to assess statistical evidence for each additional model component. For each comparison, let RSS_*r*_ and RSS_*f*_ denote the residual sums of squares of the reduced and full models, respectively, and let edf_*r*_ and edf_*f*_ denote their effective degrees of freedom. The F-type statistic is 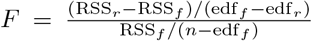. Because the spatial terms were estimated using penalized smooths, model com-plexity was represented by effective rather than nominal degrees of freedom, yielding approximate F-type tests whose empirical calibration was evaluated through simulations.

For each gene, the three model comparisons address distinct hypotheses. *F*_21_ (M2 vs. M1) tests *H*_0_ : *β*_1_ = 0, corresponding to no slide-wide H&E effect. *F*_32_ (M3 vs. M2) tests *H*_0_ : *β*_1_(**z**) ≡0, corresponding to no additional location-specific H&E effect. *F*_43_ (M4 vs. M3) tests *H*_0_ : *β*_0_(**z**) ≡*β*_0_, corresponding to no additional spatial variation beyond the total H&E effect. Among the three model-comparison statistics, *F*_32_ and *F*_43_ directly characterize spatial heterogeneity. Therefore, in the gene prioritization and pathway enrichment analysis, we focused on *F*_32_ and *F*_43_.

We summarized the sequential improvements in model fit using adjusted *R*^2^. For model M*k*, adjusted *R*^2^ was calculated as 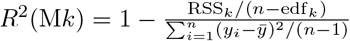, where 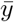 is the mean expression across locations and *R*^2^(M1) = 0.

We defined 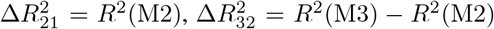, and 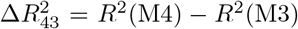. These increments correspond to adding the slide-wide H&E effect, location-specific H&E effect, and additional spatial variation, respectively. They depend on the specified model order and do not represent an order-invariant partition of gene-expression variance. We defined the remaining adjusted-*R*^2^ complement as 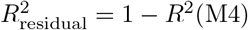.

### Gene prioritization and pathway enrichment

To prioritize genes with strong location-specific H&E effects, we retained those with 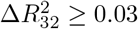 0.03 and ranked them by *F*_32_. Similarly, to prioritize genes with strong spatial variation beyond H&E, we retained those with 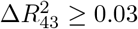 0.03 and ranked them by *F*_43_. We selected the top *N* genes from each ranking, where *N* was specified for each dataset. The two gene lists could overlap because a gene may exhibit both patterns. We performed GO Biological Process and KEGG pathway enrichment analyses separately for the two lists, with enrichment *p*-values adjusted using the BH procedure.

### Spatial interpretation of individual genes

#### Spatial component maps

For representative genes, we extracted the fitted spatial components from M4 to examine their relationships with tissue morphology. The slide-wide and location-specific H&E components were 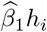 and 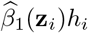, respectively. Their sum,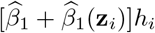, defined the total H&E effect. Spatial variation beyond H&E was represented by the centered spatial intercept,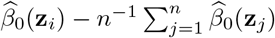.

#### Component-specific significance maps

To identify tissue regions with statistical support for the total H&E effect or spatial variation beyond H&E, we obtained pointwise estimates and standard errors of the fitted components from M4. We calculated pointwise *z*-statistics by dividing each estimate by its standard error and obtained corresponding *p*-values. For each gene and component, we applied BH adjustment across spatial locations and identified locations with adjusted *p* ≤ 0.05.

#### Correlations with cell-type proportions

To examine the relationships between the fitted spatial components and tissue cell composition, we calculated Spearman correlations between the total H&E effect or spatial variation beyond H&E and estimated cell-type proportions across locations. For single-cell-resolution data, we averaged the fitted components within spatial bins and calculated correlations with the corresponding cell-type proportions across bins.

### Simulation study

We simulated gene expression as *y*_*i*_ =*G*_*i*_ +*L*_*i*_ +*S*_*i*_ +*ε*_*i*_, where 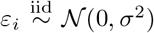, with*σ* = 1. Here,*G*_*i*_,*L*_*i*_, and*S*_*i*_ represent the simulated slide-wide H&E effect, location-specific H&E effect, and spatial variation beyond H&E, respectively. These components were generated as 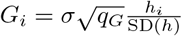,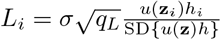, and 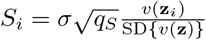.

The parameters*q*_*G*_,*q*_*L*_, and*q*_*S*_ denote component-specific signal-to-noise ratios (SNRs), defined as the variance of each simulated component divided by the noise variance. The functions*u*(**z**) and*v*(**z**) were independently generated random thin-plate regression-spline surfaces with basis dimension 10, centered and standardized across the spatial domain.

To preserve realistic spatial geometry and morphology patterns, we used the spot coordinates and H&E image from an ovarian cancer Visium dataset. The H&E feature was calculated as the mean red-channel intensity within 15× 11-pixel patches centered at each spot and mean-centered across the section. The coordinates and morphology feature were fixed across replicates, while component normalization was recalculated within each subset. Each simulation setting was evaluated over 500 replicates, with spatial surfaces and noise independently regenerated for each replicate.

The main experiment (Fig. 2) used all 3,455 spots and four configurations: {(*q*_*G*_,*q*_*L*_,*q*_*S*_) ∈ (0, 0, 0), (0.25, 0, 0), (0.25, 0.25, 0), (0.25, 0.25, 0.25)}. These settings correspond to the absence of all three modeled components, a slide-wide H&E effect only, both slide-wide and location-specific H&E effects, and all three components, respectively. We compared the distributions of*F*_32_ and*F*_43_ across these settings. For spatial recovery, we selected the median replicate among 500 based on the Pearson correlation between true and estimated spatial variation beyond H&E. Additional simulations varying sample size, component SNR, and concurvity are described in the Supplementary Information.

### Real-data analyses

**Human dorsolateral prefrontal cortex dataset**. We used log-transformed expression and retained 8,919 genes with a standard deviation greater than 0.2 across spots. The morphology covariate was the mean green-channel intensity within 7 × 7-pixel H&E patches centered at each Visium spot and mean-centered across the section. After applying the threshold of Δ*R*^2^ ≥ 0.03, we selected the top 200 genes ranked by*F*_32_ and*F*_43_, respectively. GO Biological Process and KEGG pathway enrichment analyses used clusterProfiler [30]. We obtained precomputed proportions of nine cell types from Spacelink [31] and calculated Spearman correlations with the fitted total H&E effect or spatial variation beyond H&E for the 102 genes shared by the two top-200 lists. For component-based clustering, we applied*k*-means with*k* = 7 to the corresponding fitted components for the two top-200 gene sets.

#### Xenium breast cancer

We used segmented cell centers and log-transformed expression for model fitting. We extracted hierarchical histology features using iSTAR [15] and used the first principal component (PC1) as the morphology covariate. After the same ranking procedure, we selected the top 50 genes from each ranking for GO Biological Process and KEGG pathway enrichment. Using the provided cell annotations, we grouped cells into 100 × 100*µ*m bins containing at least ten cells and calculated cell-type proportions. We averaged the fitted total H&E effect and spatial variation beyond H&E within each bin and calculated Spearman correlations with cell-type proportions for the 36 genes shared by the two top-50 lists. For the four ECM genes, stromal-rich bins contained ≥50% stromal cells, while tumour-rich bins contained ≥ 50% DCIS or invasive-tumour cells. We calculated distances from stromal-rich bins to the nearest tumour-rich bin and averaged component scores within [0, 200], (200, 300], (300, 500], and*>* 500*µ*m.

#### HER2-positive breast cancer

We analyzed sections A1, G2, and H1 separately, retaining genes detected in at least 10% of spots. Counts were log-transformed, and the morphology covariate was the mean RGB intensity within approximately 100*µ*m-wide H&E patches centered at each spot and mean-centered within each section. We identified invasive-cancer markers using two-sided Wilcoxon rank-sum tests comparing invasive-cancer with other annotated regions, retaining genes with positive*z*-statistics, higher mean expression, and BH-adjusted*p <* 0.05 in each section, yielding 25 shared markers. We then calculated Spearman correlations between proportions of eight major cell types from HER2ST [28] and the fitted total H&E effect or spatial variation beyond H&E for these markers.

## Data and code availability

All datasets used in this study are publicly available. The main real-data analyses used three datasets: (1) the human DLPFC Visium dataset (sample 151673) from spatialLIBD [32, 25]; (2) the Xenium human breast cancer dataset from 10x Genomics [6]; and (3) the HER2ST dataset from Zenodo [28]. The analyses in the Supplementary Information used two additional datasets: (4) the human ovarian cancer Visium FFPE dataset from 10x Genomics [33], also used for simulations; and (5) the bladder cancer data from the MorphLink repository [24]. MorphoANOVA has been implemented as an R package, which is available at https://github.com/powei219/MorphoANOVA.

## Notes

https://github.com/powei219/MorphoANOVA

### Competing Interest Statement

The authors have declared no competing interest.

## References

[1] Patrik L Ståhl, Fredrik Salmén, Sanja Vickovic, Anna Lundmark, José Fernández Navarro, Jens Magnusson, Stefania Giacomello, Michaela Asp, Jakub O Westholm, Mikael Huss, et al. Visualization and analysis of gene expression in tissue sections by spatial transcriptomics. Science, 353(6294):78–82, 2016.

[2] Samuel G Rodriques, Robert R Stickels, Aleksandrina Goeva, Carly A Martin, Evan Murray, Charles R Vanderburg, Joshua Welch, Linlin M Chen, Fei Chen, and Evan Z Macosko. Slide-seq: A scalable technology for measuring genome-wide expression at high spatial resolution. Science, 363(6434):1463–1467, 2019.

[3] Sanja Vickovic, Gö kcen Eraslan, Fredrik Salmén, Johanna Klughammer, Linnea Stenbeck, Denis Schapiro, Tarmo Äijö, Richard Bonneau, Ludvig Bergenstråhle, José Fernandéz Navarro, et al. High-definition spatial transcriptomics for in situ tissue profiling. Nature Methods, 16(10):987–990, 2019.

[4] Kok Hao Chen, Alistair N Boettiger, Jeffrey R Moffitt, Siyuan Wang, and Xiaowei Zhuang. Spatially resolved, highly multiplexed RNA profiling in single cells. Science, 348(6233):aaa6090, 2015.

[5] Sheel Shah, Eric Lubeck, Wen Zhou, and Long Cai. seqFISH accurately detects transcripts in single cells and reveals robust spatial organization in the hippocampus. Neuron, 94(4):752–758, 2017.

[6] 10x Genomics. Xenium in situ human breast dataset explorer, 2024. Accessed: 2024-07-13.

[7] Jordan T Ash, Gregory Darnell, Daniel Munro, and Barbara E Engelhardt. Joint analysis of expression levels and histological images identifies genes associated with tissue morphology. Nature Communications, 12(1):1609, 2021.

[8] Ankit Agrawal, Stefan Thomann, Sukanya Basu, and Dominic Grön. NiCo identifies extrinsic drivers of cell state modulation by niche covariation analysis. Nature Communications, 15(1):10628, 2024.

[9] David Gosselin, Verena M Link, Casey E Romanoski, Gregory J Fonseca, Dawn Z Eichenfield, Nathanael J Spann, Joshua D Stender, Hyun B Chun, Hannah Garner, Frederic Geissmann, et al. Environment drives selection and function of enhancers controlling tissue-specific macrophage identities. Cell, 159(6):1327–1340, 2014.

[10] Rasa Elmentaite, Cecilia Domínguez Conde, Lu Yang, and Sarah A Teichmann. Single-cell atlases: shared and tissue-specific cell types across human organs. Nature Reviews Genetics, 23(7):395–410, 2022.

[11] Eduard Chelebian, Christophe Avenel, and Carolina Wählby. Combining spatial transcriptomics with tissue morphology. Nature Communications, 16(1):4452, 2025.

[12] Bryan He, Ludvig Bergenstråhle, Linnea Stenbeck, Abubakar Abid, Alma Andersson, Åke Borg, Jonas Maaskola, Joakim Lundeberg, and James Zou. Integrating spatial gene expression and breast tumour morphology via deep learning. Nature Biomedical Engineering, 4(8):827–834, 2020.

[13] Yuansong Zeng, Zhuoyi Wei, Weijiang Yu, Rui Yin, Yuchen Yuan, Bingling Li, Zhonghui Tang, Yutong Lu, and Yuedong Yang. Spatial transcriptomics prediction from histology jointly through transformer and graph neural networks. Briefings in Bioinformatics, 23(5), 2022.

[14] Ronald Xie, Kuan Pang, Sai Chung, Catia Perciani, Sonya MacParland, Bo Wang, and Gary Bader. Spatially resolved gene expression prediction from histology images via bi-modal contrastive learning. Advances in Neural Information Processing Systems, 36:70626–70637, 2023.

[15] Daiwei Zhang, Amelia Schroeder, Hanying Yan, Haochen Yang, Jian Hu, Michelle YY Lee, Kyung S Cho, Katalin Susztak, George X Xu, Michael D Feldman, et al. Inferring super-resolution tissue architecture by integrating spatial transcriptomics with histology. Nature Biotechnology, 42(9):1372–1377, 2024.

[16] Xiao Tan, Andrew Su, Minh Tran, and Quan Nguyen. SpaCell: integrating tissue morphology and spatial gene expression to predict disease cells. Bioinformatics, 36(7):2293–2294, 2020.

[17] Duy Pham, Xiao Tan, Brad Balderson, Jun Xu, Laura F Grice, Sohye Yoon, Emily F Willis, Minh Tran, Pui Yeng Lam, Arti Raghubar, et al. Robust mapping of spatiotemporal trajectories and cell–cell interactions in healthy and diseased tissues. Nature Communications, 14(1):7739, 2023.

[18] Kaiming He, Xiangyu Zhang, Shaoqing Ren, and Jian Sun. Deep residual learning for image recognition. In Proceedings of the IEEE Conference on Computer Vision and Pattern Recognition, pages 770–778, 2016.

[19] Jian Hu, Xiangjie Li, Kyle Coleman, Amelia Schroeder, Nan Ma, David J Irwin, Edward B Lee, Russell T Shinohara, and Mingyao Li. SpaGCN: Integrating gene expression, spatial location and histology to identify spatial domains and spatially variable genes by graph convolutional network. Nature Methods, 18(11):1342–1351, 2021.

[20] Yongshuo Zong, Tingyang Yu, Xuesong Wang, Yixuan Wang, Zhihang Hu, and Yu Li. conST: an interpretable multi-modal contrastive learning framework for spatial transcriptomics. bioRxiv, page 2022.01.14.476408, 2022.

[21] Yuansong Zeng, Rui Yin, Mai Luo, Jianing Chen, Zixiang Pan, Yutong Lu, Weijiang Yu, and Yuedong Yang. Identifying spatial domain by adapting transcriptomics with histology through contrastive learning. Briefings in Bioinformatics, 24(2):bbad048, 2023.

[22] Liviu Badea and Emil Stănescu. Identifying transcriptomic correlates of histology using deep learning. PLOS ONE, 15(11):e0242858, 2020.

[23] Sungwoo Bae, Hongyoon Choi, and Dong Soo Lee. Discovery of molecular features underlying the morphological landscape by integrating spatial transcriptomic data with deep features of tissue images. Nucleic Acids Research, 49(10):e55, 2021.

[24] Jing Huang, Chenyang Yuan, Jiahui Jiang, Jianfeng Chen, Sunil S Badve, Yesim Gokmen-Polar, Rossana L Segura, Xinmiao Yan, Alexander Lazar, Jianjun Gao, et al. Bridging cell morphological behaviors and molecular dynamics in multi-modal spatial omics with MorphLink. Nature Communications, 16(1):5878, 2025.

[25] Kristen R Maynard, Leonardo Collado-Torres, Lukas M Weber, Cedric Uytingco, Brianna K Barry, Stephen R Williams, Joseph L Catallini, Matthew N Tran, Zachary Besich, Madhavi Tippani, et al. Transcriptome-scale spatial gene expression in the human dorsolateral prefrontal cortex. Nature Neuroscience, 24(3):425–436, 2021.

[26] Wendy Xin, Megumi Kaneko, Richard H. Roth, Albert Zhang, Sonia Nocera, Jun B. Ding, Michael P. Stryker, and Jonah R. Chan. Oligodendrocytes and myelin limit neuronal plasticity in visual cortex. Nature, 633:856–863, 2024.

[27] Ran Wang, Yan Qian, Xiaojing Guo, Fangda Song, Zhiqiang Xiong, Shirong Cai, Xiuwu Bian, Man Hon Wong, Qin Cao, Lixin Cheng, et al. STModule: identifying tissue modules to uncover spatial components and characteristics of transcriptomic landscapes. Genome Medicine, 17(1):18, 2025.

[28] Alma Andersson, Ludvig Larsson, Linnea Stenbeck, Fredrik Salmén, Anna Ehinger, Sunny Z. Wu, Ghamdan Al-Eryani, Daniel Roden, Alex Swarbrick, Åke Borg, Jonas Frisén, Camilla Engblom, and Joakim Lundeberg. Spatial deconvolution of HER2-positive breast cancer delineates tumor-associated cell type interactions. Nature Communications, 12(1):6012, 2021.

[29] Shan Yu and Wei Vivian Li. spVC for the detection and interpretation of spatial gene expression variation. Genome Biology, 25(1):103, 2024.

[30] Tianzhi Wu, Erqiang Hu, Shuangbin Xu, Meijun Chen, Pingfan Guo, Zehan Dai, Tingze Feng, Lang Zhou, Wenli Tang, Li Zhan, Xiaocong Fu, Shanshan Liu, Xiaochen Bo, and Guangchuang Yu. clusterProfiler 4.0: A universal enrichment tool for interpreting omics data. The Innovation, 2(3):100141, 2021.

[31] Hanbyul Lee, Haochen Sun, Xuewei Cao, Berke Karaahmet, Zhijian Li, Hans Ulrich-Klein, Mariko Taga, Gao Wang, Philip L. De Jager, David A. Bennett, Luca Pinello, Xin Jin, Rahul Mazumder, and Kushal K. Dey. Mapping disease critical spatially variable gene programs by integrating spatial transcriptomics with human genetics. bioRxiv, 2025. Preprint.

[32] Brenda Pardo, Abby Spangler, Lukas M Weber, Stephanie C Page, Stephanie C Hicks, Andrew E Jaffe, Keri Martinowich, Kristen R Maynard, and Leonardo Collado-Torres. spatialLIBD: an R/Bioconductor package to visualize spatially-resolved transcriptomics data. BMC Genomics, 23(1):434, 2022.

[33] 10x Genomics. Visium FFPE human ovarian cancer web summary. https://cf.10xgenomics.com/samples/spatial-exp/1.3.0/Visium_FFPE_Human_Ovarian_Cancer/Visium_FFPE_Human_Ovarian_Cancer_web_summary.html, 2024. Accessed: 2024-07-04.

