## Supplementary Information for "MorphoANOVA: Dissecting Histology–Gene Expression Relationships in Spatial Transcriptomics"

#### Supplementary Figures

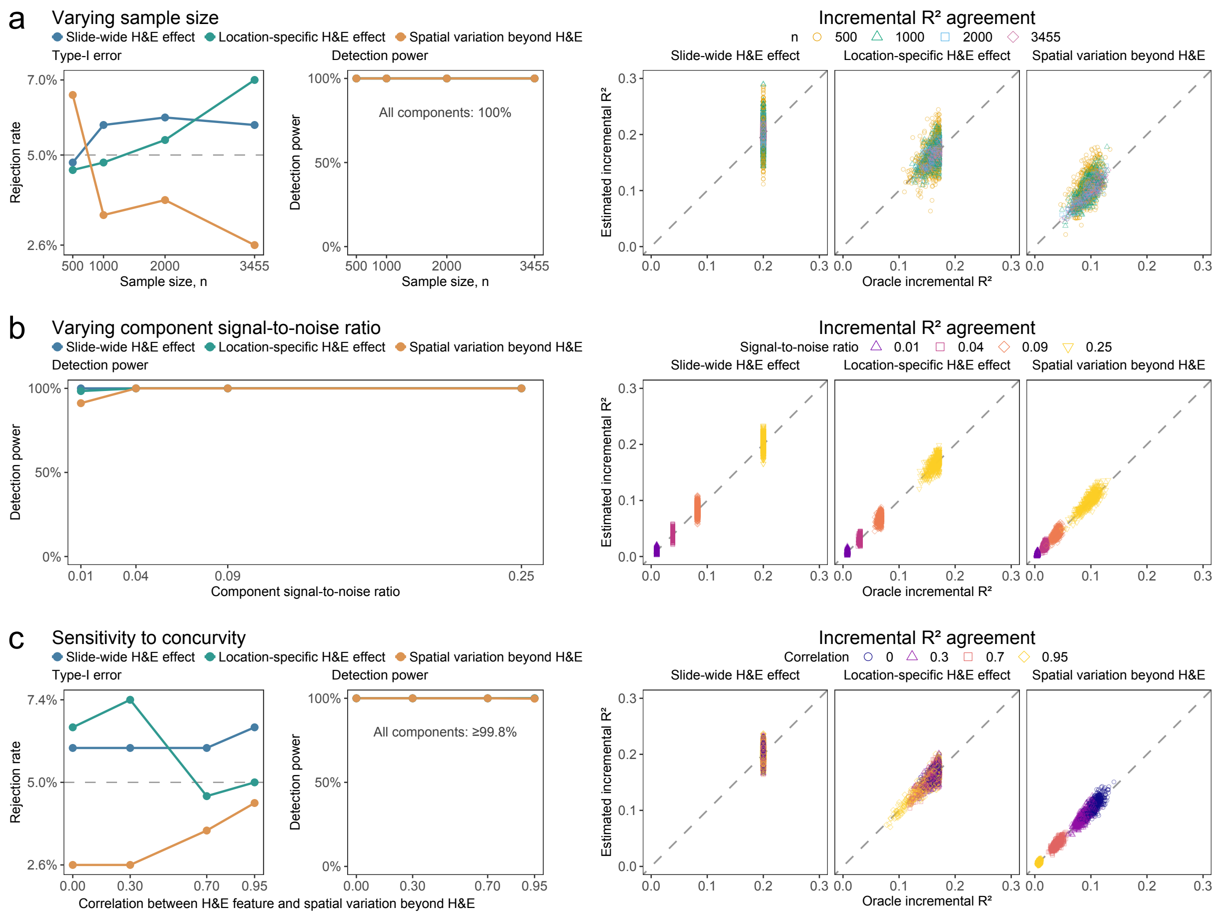


Supplementary Figure 1: **MorphoANOVA inference across simulation settings.** (**a**) Type-I error, detection power and incremental $R^2$ agreement across $n = 500, 1000, 2000$ and 3455 spatial locations. (**b**) Detection power and incremental $R^2$ agreement at four nonzero component signal-to-noise ratios (SNRs: $0.01, 0.04, 0.09, 0.25$). (**c**) Type-I error, detection power and incremental $R^2$ agreement across correlations of $0, 0.3, 0.7$ and 0.95 between a synthetic H&E feature and spatial variation beyond H&E (concurvity). Panels (**b**,**c**) used $n = 3455$. Power and agreement analyses in (**a**,**c**) used a component SNR of 0.25. Each setting included 500 replicates.

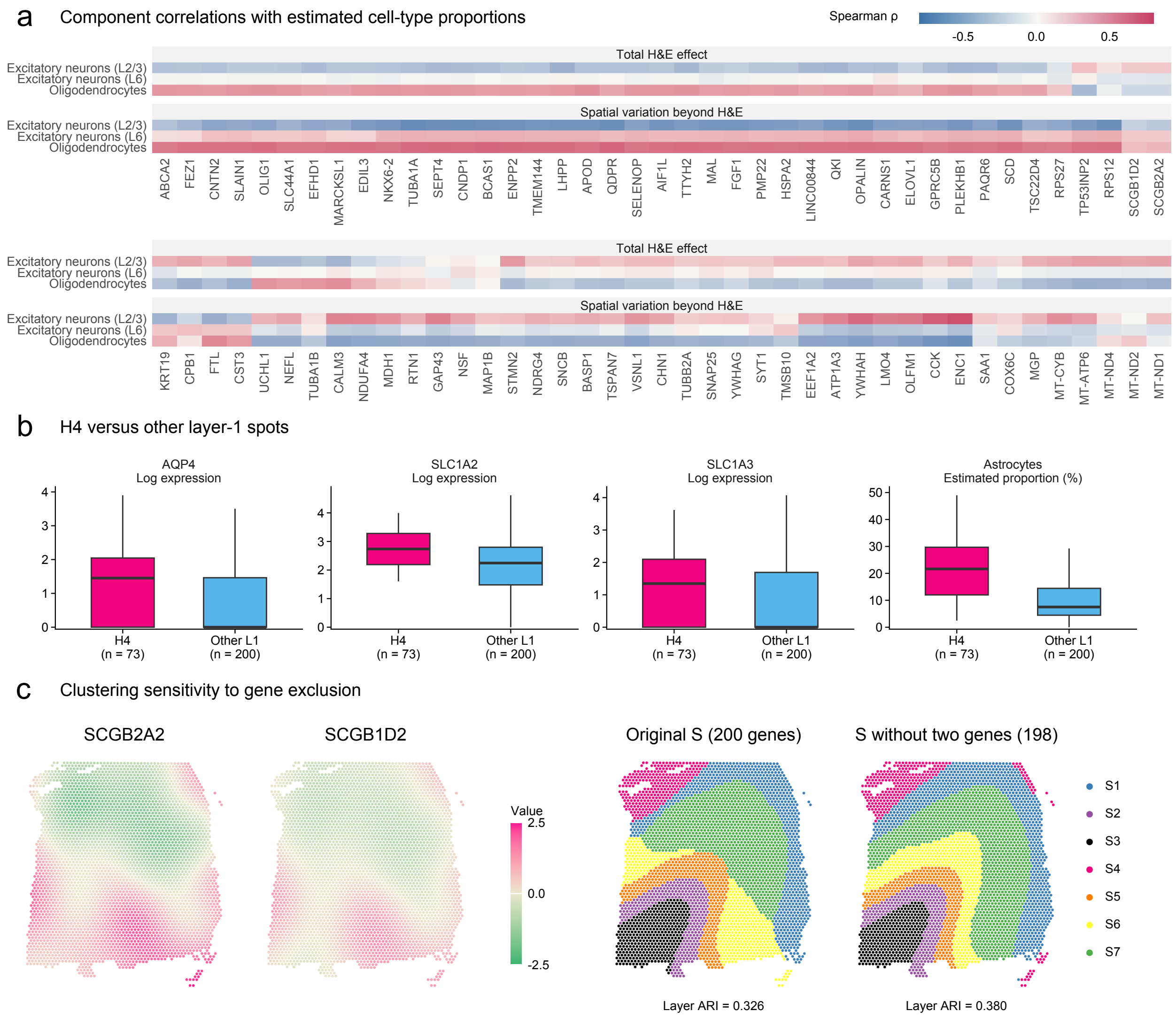


Supplementary Figure 2: **Additional gene expression and functional enrichment in the human cortex.** (**a**) Spatial expression maps of the top six genes from each $F_{32}$- and $F_{43}$-based ranking. (**b**) GO Biological Process and KEGG pathway enrichment of the corresponding top-200 lists. Dot size and colour indicate gene count and BH-adjusted $p$-value, respectively.

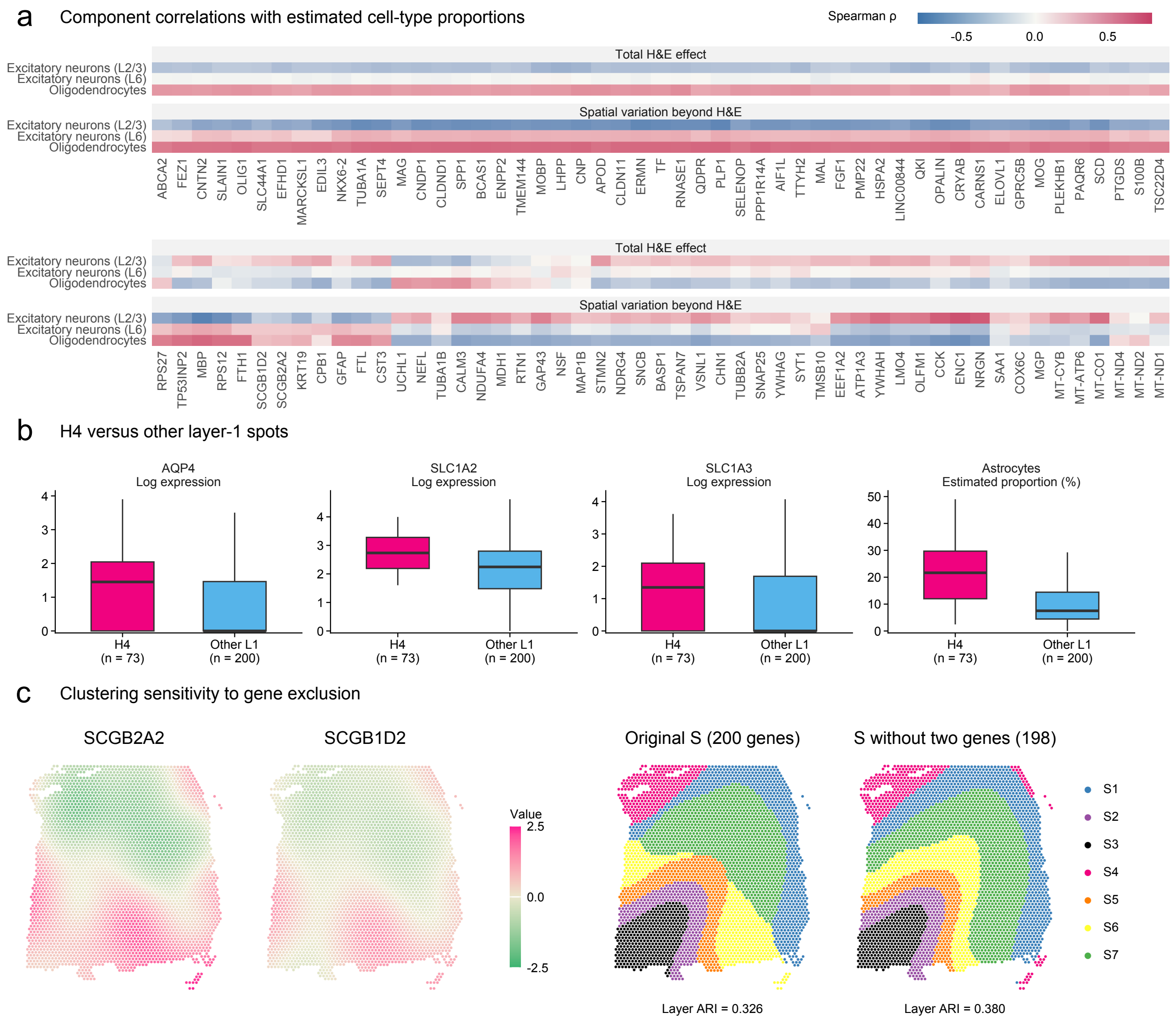


Supplementary Figure 3: **Cellular associations and clustering of cortical expression components.** (**a**) Spearman correlations of the total H&E effect and spatial variation beyond H&E with estimated cell-type proportions across spots for the remaining 82 of the 102 genes shared by the two top-200 lists. (**b**) Observed expression of three astrocyte markers (*AQP4*, *SLC1A2*, and *SLC1A3*) and estimated astrocyte proportions in H4 and the remaining L1 spots. Boxes show medians and interquartile ranges. (**c**) Fitted spatial variation beyond H&E for *SCGB2A2* and *SCGB1D2*, and clustering based on this component before and after excluding these two genes from the $F_{43}$-based top-200 list. ARI indicates agreement with anatomical annotations.

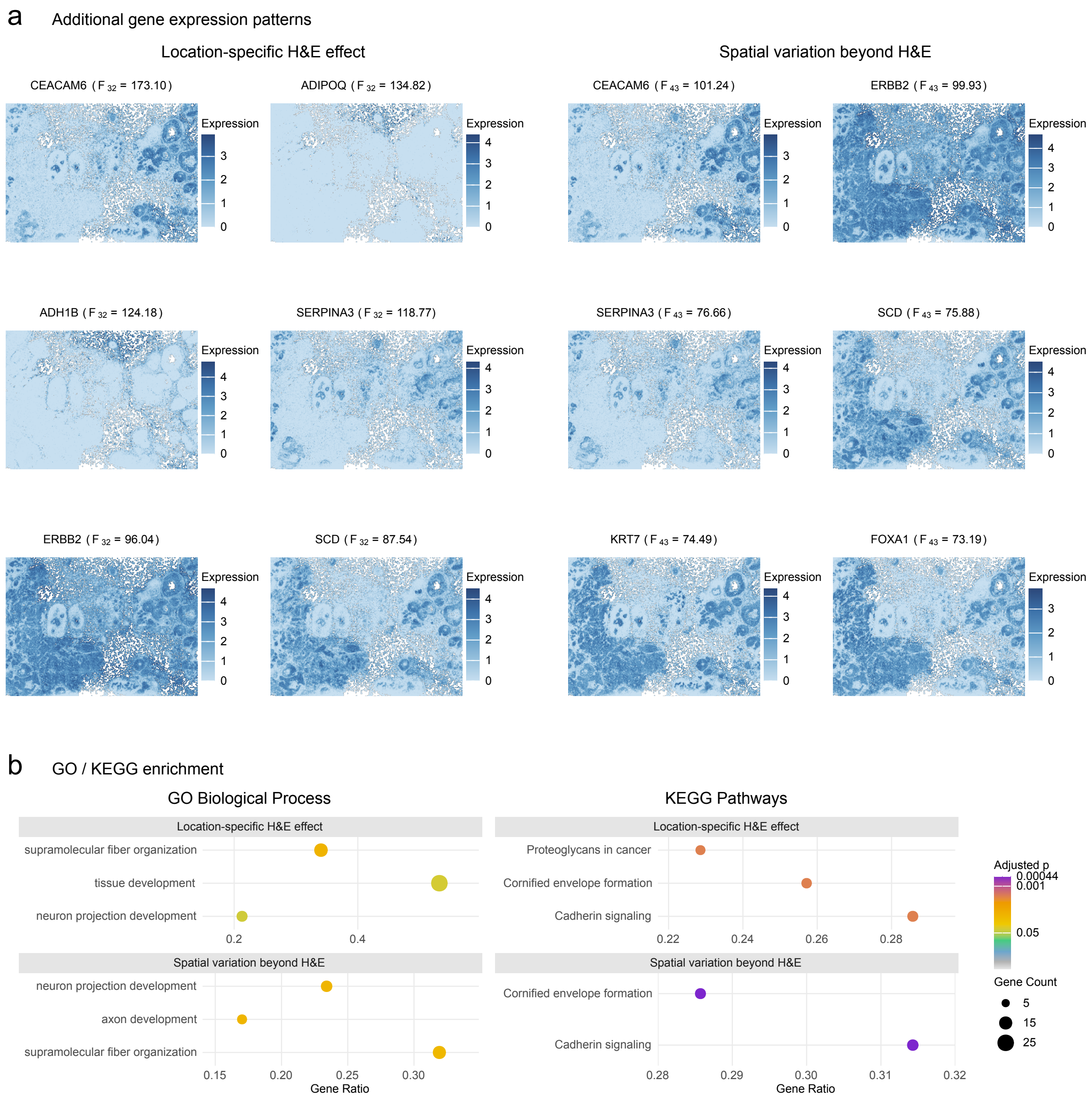


Supplementary Figure 4: **Additional gene expression and functional enrichment in breast cancer.** (**a**) Spatial expression maps of the top six genes from each $F_{32}$- and $F_{43}$-based ranking. (**b**) GO Biological Process and KEGG pathway enrichment of the corresponding top-50 lists. Dot size and colour indicate gene count and BH-adjusted $p$-value, respectively.

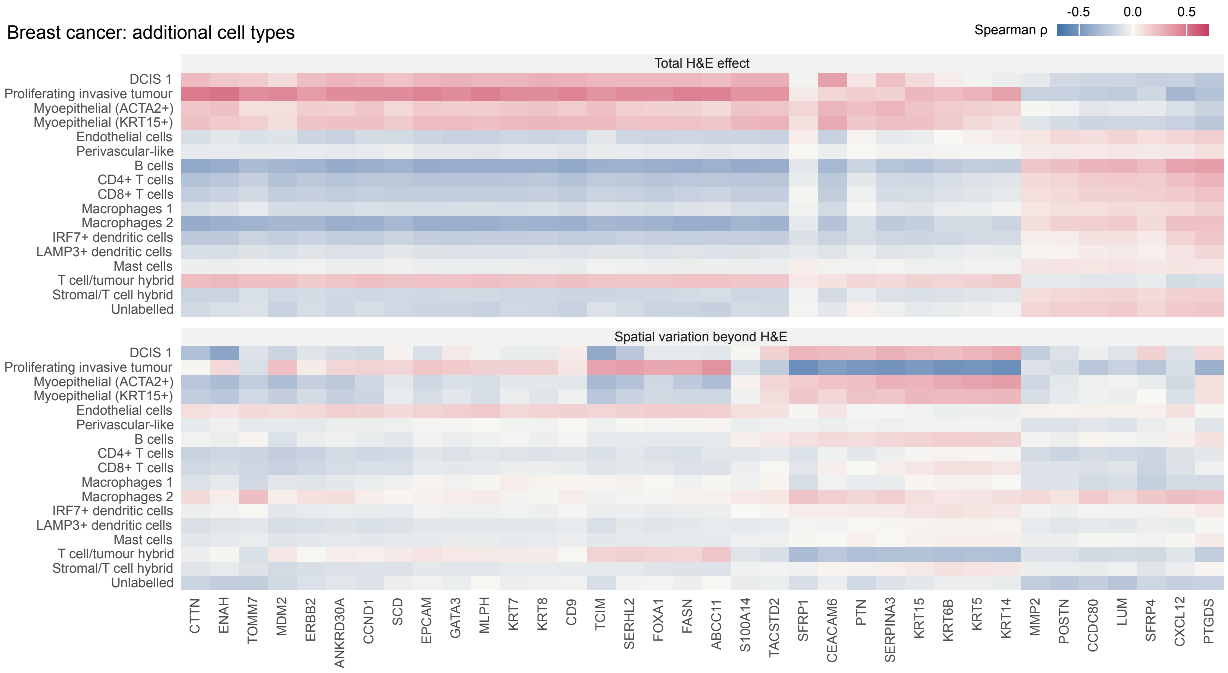


Supplementary Figure 5: **Additional cellular associations in breast cancer.** Spearman correlations of the total H&E effect and spatial variation beyond H&E with proportions of 17 additional cell categories for the 36 genes shared by the two top-50 lists. Correlations were calculated across 100 $\mu$m spatial bins.

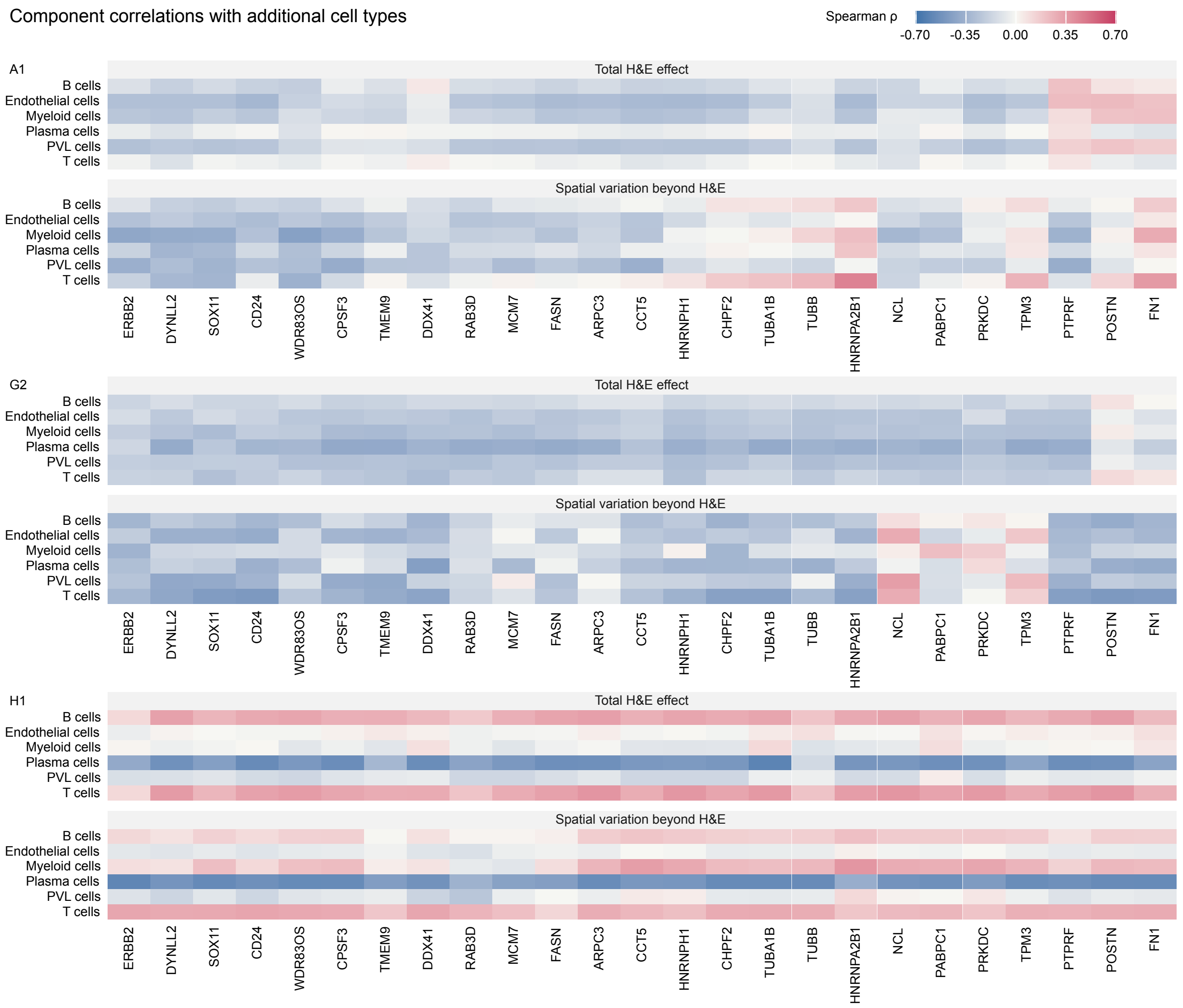


Supplementary Figure 6: **Cellular associations with six additional cell types in HER2$^+$ breast cancer.** Spearman correlations of the total H&E effect and spatial variation beyond H&E with estimated cell-type proportions across all spots in sections A1, G2, and H1.

### Supplementary Results

#### Distinct morphogenomic patterns in the ovarian tumour microenvironment

We investigated how expression patterns relate to tissue morphology in ovarian cancer, and where additional spatial expression patterns remain beyond the selected H&E feature. The dataset comprised measurements of 17,294 genes across 3,455 spots [1], with 2,977 genes retained after expression-variability filtering. Local histology was represented by a mean-centered red-channel intensity feature extracted at each spot (Supplementary Fig. 7a).

Genes with location-specific H&E effects and spatial variation beyond H&E shared antigen-presentation enrichment but differed in other biological processes. Following the component-specific 3% variance-explained filter, we ranked genes by $F_{32}$ and $F_{43}$ and selected the top 100 from each ranking (Supplementary Fig. 7b; top six shown in Supplementary Fig. 8). GO enrichment for location-specific H&E effects included antigen processing and presentation and MHC class II complex assembly, whereas spatial variation beyond H&E was associated with extracellular matrix organization and humoral immune responses. KEGG analysis showed enrichment for antigen processing and presentation in both lists. The location-specific H&E list was also enriched for herpes simplex virus 1 infection and phagocytosis, whereas the spatial variation beyond H&E list included ECM–receptor interaction and cytoskeleton in muscle cells (Supplementary Fig. 7c).

We selected *IGHM*, which ranked highly in both lists, to examine its morphogenomic pattern. The fitted model explained 80.0% of its expression variation, including 18.4% attributable to the slide-wide H&E effect, 20.0% to the location-specific H&E effect, and 41.6% to spatial variation beyond H&E (Supplementary Fig. 7d). Notably, the significance maps showed that the total H&E effect was concentrated along a narrow, curved band traversing the tissue interior, whereas spatial variation beyond H&E extended across much broader regions (Supplementary Fig. 7e). Spearman correlations with cell proportions showed associations with lymphocytes for both components ($\rho = 0.34$ and 0.41, respectively). Spatial variation beyond H&E showed stronger associations with stromal-cell ($\rho = -0.51$ versus $-0.05$) and red-blood-cell proportions ($\rho = 0.41$ versus $-0.08$; Supplementary Fig. 7f).

The two components showed distinct associations with the cellular composition of the ovarian tumour microenvironment. Across the 59 genes shared by the two top-100 lists, spatial variation beyond H&E generally showed stronger absolute correlations with lymphocyte, stromal-cell and red-blood-cell proportions, whereas the total H&E effect was more strongly associated with tumour-cell proportions (Supplementary Fig. 7g). Correlations with macrophage and necrosis/karyorrhexis proportions are shown in Supplementary Fig. 8.

#### Spatial interpretation of morphology–gene associations identified by MorphLink

To assess whether MorphoANOVA could provide a spatial interpretation of morphology–gene associations identified by an external image-analysis framework, we analyzed a human bladder cancer Visium dataset previously studied using MorphLink [2]. MorphLink identified the interquartile range (IQR) of nuclear solidity as a morphology feature associated with antigen-presentation genes using the Curve-based Pattern Similarity Index (CPSI), which summarizes morphology–gene concordance across the tissue. We used the same nuclear-solidity feature as the morphology covariate in MorphoANOVA (Supplementary Fig. 9a), enabling direct comparison of gene prioritization while further examining how these associations varied across tissue regions.

Across the antigen-presentation gene set, gene ranks based on the MorphoANOVA location-specific H&E statistic $F_{32}$ were strongly concordant with those based on CPSI (Spearman's $\rho = 0.849$; Supplementary Fig. 9b). This agreement showed that MorphoANOVA recovered the morphology-associated transcriptional signal identified by MorphLink.

MorphoANOVA further resolved the spatial variation underlying these morphology–gene associations. For *B2M*, the top-ranked gene by $F_{32}$, the slide-wide H&E effect explained 7.9% of expression variation, whereas the location-specific H&E effect explained an additional 48.4% (Supplementary Fig. 9c). The component maps showed regional structure beyond the slide-wide association. Similar decompositions were observed for *CALR*, *CD74*, and *TAP1*, with location-specific H&E effects explaining substantially more variation than slide-wide H&E effects (Supplementary Fig. 10). Thus, MorphoANOVA extended CPSI-based pattern-similarity analysis by localizing spatially heterogeneous morphology-associated variation and quantifying their contributions relative to the slide-wide association.

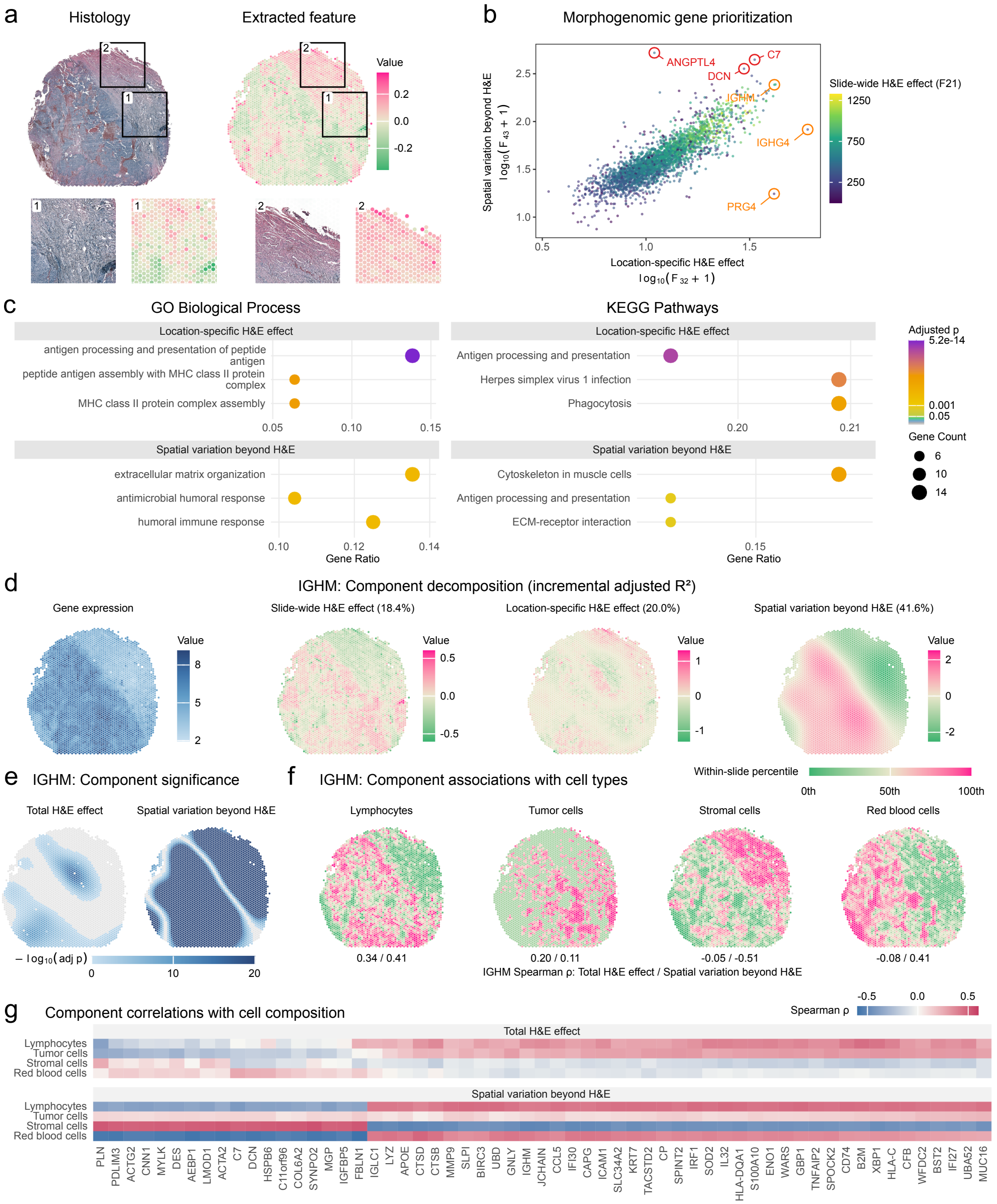


Supplementary Figure 7: **Morphogenomic patterns in human ovarian cancer.** (**a**) H&E image and extracted morphology feature, with enlarged views of two marked regions. (**b**) Morphogenomic gene prioritization of $\log_{10}(F_{32}+1)$ and $\log_{10}(F_{43}+1)$, colored by $F_{21}$. Orange and red labels mark the top three genes in the respective rankings. (**c**) GO Biological Process and KEGG pathway enrichment of the two top-100 lists. Dot size and color indicate gene count and BH-adjusted $p$-value, respectively. (**d**) Observed *IGHM* expression and three fitted components. Percentages indicate sequential contributions to expression variation. (**e**) BH-adjusted pointwise $p$-value maps for the total H&E effect and spatial variation beyond H&E. (**f**) Image-derived cell-category proportions shown as within-slide percentiles. Paired values give Spearman correlations with the *IGHM* total H&E effect and spatial variation beyond H&E, respectively. (**g**) Corresponding correlations for the 59 genes shared by the two top-100 lists.

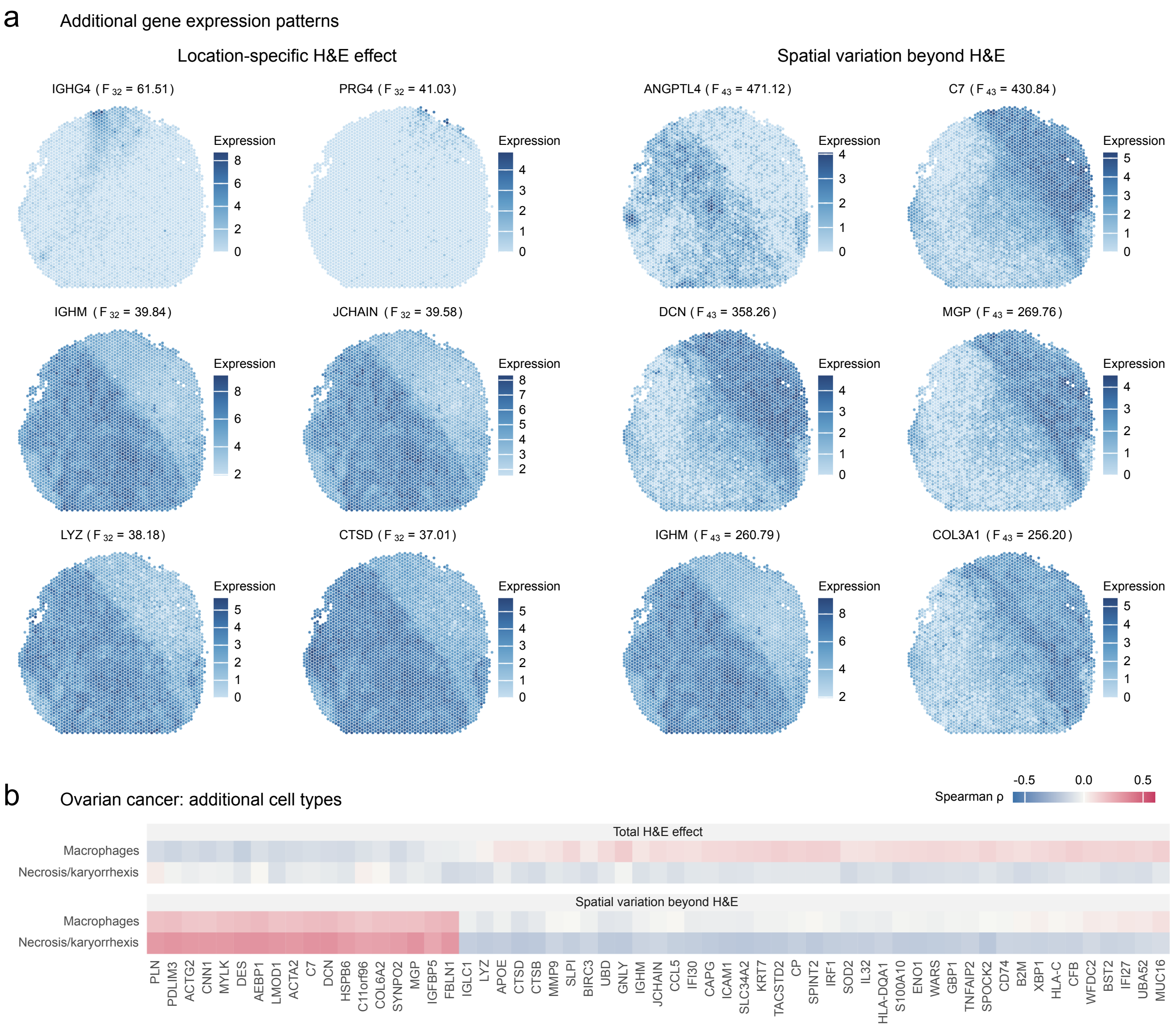


Supplementary Figure 8: **Additional spatial expression patterns and cellular associations in ovarian cancer. (a)** Spatial expression maps of the top six genes from each of the $F_{32}$- and $F_{43}$-based rankings. Labels indicate the corresponding $F$-statistics. **(b)** Spearman correlations of the fitted total H&E effect and spatial variation beyond H&E with macrophage and necrosis/karyorrhexis proportions for the 59 genes shared by the two top-100 lists.

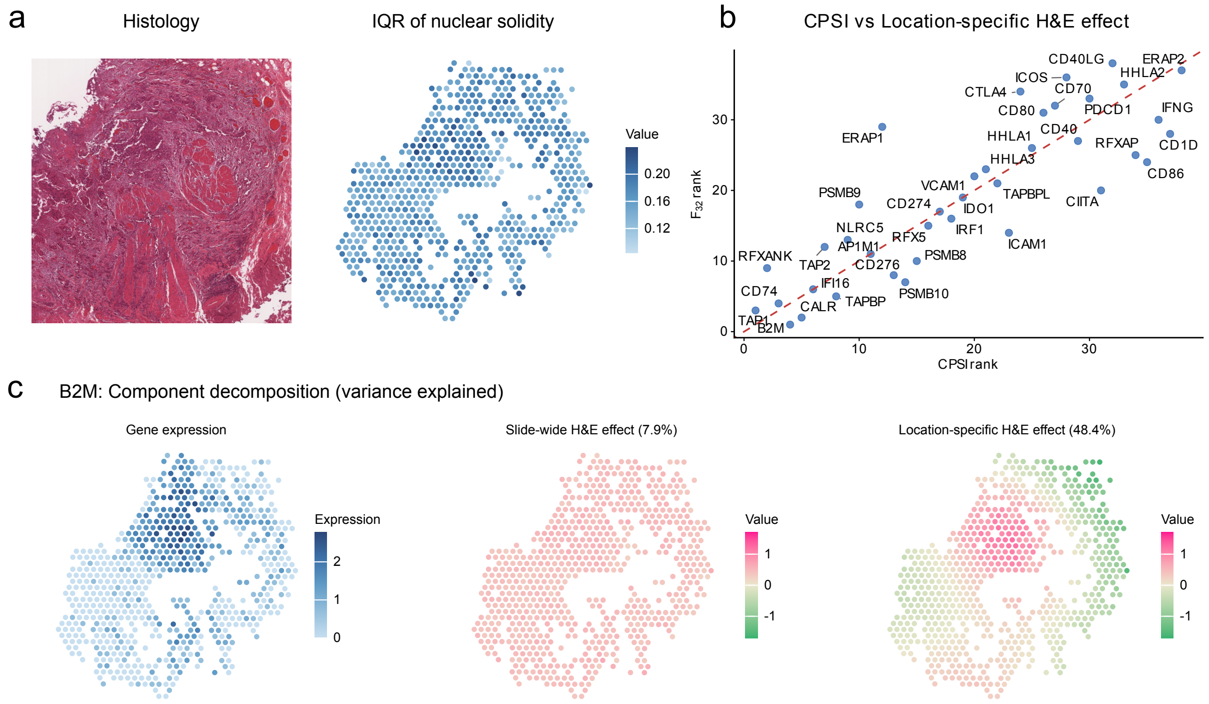


Supplementary Figure 9: **Spatial interpretation of morphology–gene associations identified by MorphLink.** (**a**) H&E image and nuclear-solidity IQR in human bladder cancer. (**b**) CPSI and $F_{32}$ ranks for the original 38 genes. (**c**) Observed *B2M* expression and its slide-wide and location-specific H&E effects. Percentages indicate sequential contributions to expression variation.

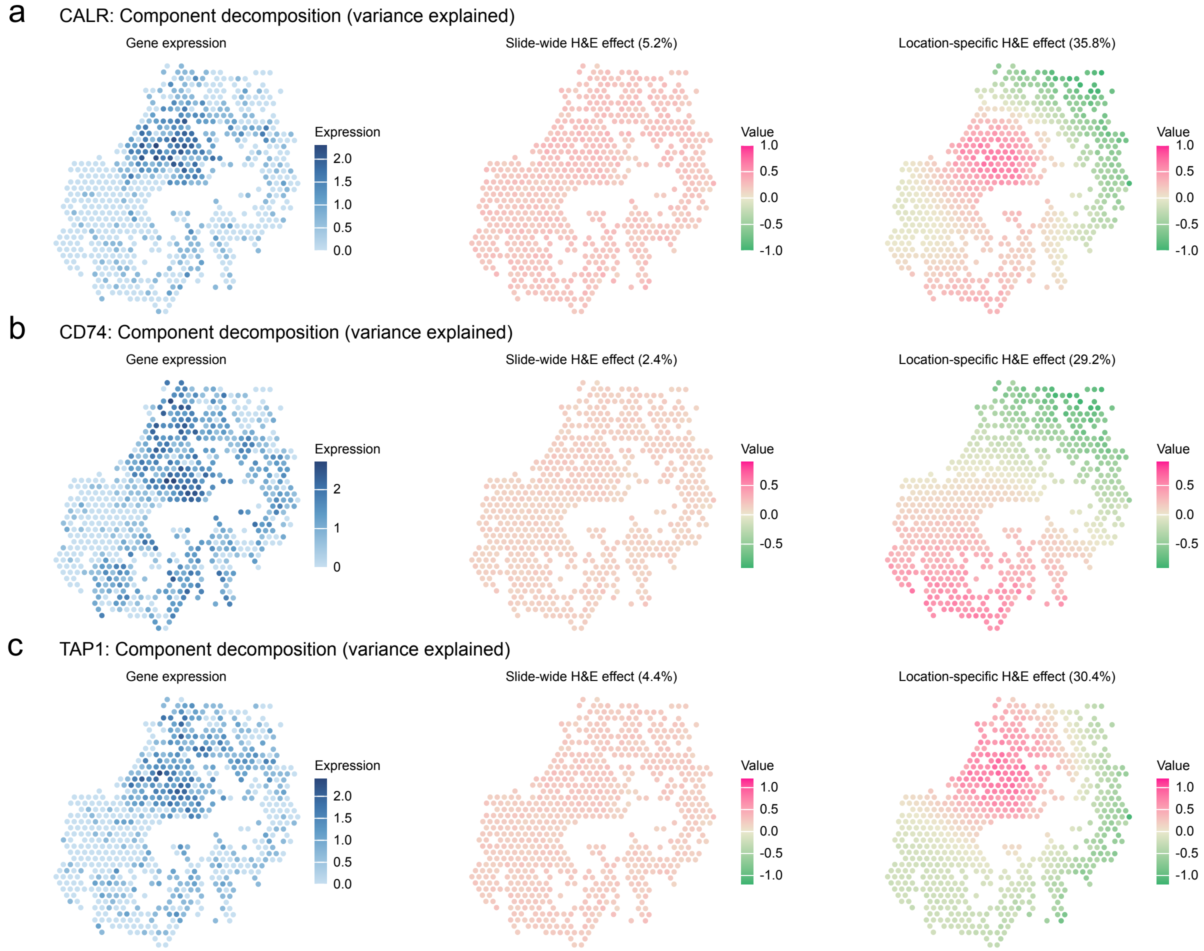


Supplementary Figure 10: **Additional spatial decompositions of MorphLink-associated genes.** Observed expression and fitted slide-wide and location-specific H&E effects for (**a**) *CALR*, (**b**) *CD74*, and (**c**) *TAP1*. Percentages indicate sequential contributions to expression variation.

### Supplementary Methods

#### Additional simulation settings

Additional simulations evaluated sample size, component SNR, and concurvity separately (Supplementary Fig. 1). Sample-size analyses used nested maximin subsets with $n \in \{500, 1000, 2000, 3455\}$. Component-SNR analyses varied the tested component's SNR $q \in \{0.01, 0.04, 0.09, 0.25\}$ at $n = 3455$. Both analyses used the observed H&E feature. For each model comparison, preceding components remained at an SNR of 0.25 and subsequent components were absent.

Concurvity analyses used synthetic features $h^{(\rho)} = \rho U + \sqrt{1-\rho^2} Z$, where $U$ is the centered, standardized realization of $v(\mathbf{z})$, and $Z$ is the observed H&E feature after centering, removing its projection onto $U$, and standardization. We varied $\rho \in \{0, 0.3, 0.7, 0.95\}$ at $n = 3455$. Power and incremental adjusted $R^2$ analyses in the sample-size and concurvity settings used a tested-component SNR of 0.25. Type-I error was the proportion of $p$-values below 0.05 with the tested component absent, and detection power was the rejection proportion with that component present. Each setting contained 500 replicates.

To evaluate incremental adjusted $R^2$, we constructed signal-based oracle references from the known conditional mean $\boldsymbol{\mu} = \mathbf{G} + \mathbf{L} + \mathbf{S}$. Let $P_k$ denote the unpenalized orthogonal projector onto the design space of model M$k$, constructed using the same spatial spline bases as the fitted models. We calculated $R^2_{k,\text{oracle}} = 1 - \frac{\|\boldsymbol{\mu} - P_k \boldsymbol{\mu}\|^2 + (n-1)\sigma^2}{\|\boldsymbol{\mu} - \bar{\mu}\mathbf{1}\|^2 + (n-1)\sigma^2}$, with successive differences defining the three oracle contributions. These references quantify contributions within the model spaces and were compared with the corresponding MorphoANOVA estimates at nonzero tested-component SNRs.

#### Additional details of the ovarian cancer analysis and MorphLink comparison

For the human ovarian cancer Visium dataset, we retained genes with a raw-count standard deviation greater than 2 across spots, resulting in 2,977 genes for analysis. The retained counts were transformed using log(count + 1) before model fitting. To construct the morphology covariate, we calculated the mean red-channel intensity within a $15 \times 11$-pixel image patch centered at each Visium spot. MorphoANOVA was then fitted separately to each retained gene. The spatial spline basis dimension was 30. For downstream analysis, we applied a component-specific incremental adjusted $R^2$ threshold of 0.03 and selected the top 100 eligible genes ranked by $F_{32}$ and $F_{43}$, respectively. GO Biological Process and human KEGG pathway enrichment analyses were performed using the R package `clusterProfiler` [3], using all analyzed genes as the background universe. Enrichment $p$-values were adjusted using the BH procedure, and terms with BH-adjusted $p < 0.05$ were considered significant. Redundant GO terms were reduced using Wang semantic similarity with a cutoff of 0.7, retaining the term with the smallest adjusted $p$-value. For region-level interpretation, we selected *IGHM* as a representative gene. For each of the 59 genes shared by the two top-100 lists, we calculated Spearman correlations across spots between cell-type proportions and the fitted total H&E effect or spatial variation beyond H&E. Cell-type proportions were calculated from the image-derived cell counts provided with the iIMPACT study [4].

For the human bladder cancer Visium dataset used in the MorphLink comparison, we followed the original MorphLink preprocessing. After transforming the raw counts using log(count + 1), we analyzed 665 spots and the 38 genes available from the antigen-presentation gene set reported in Supplementary Table 5 of the original MorphLink study [2]. To facilitate a fair comparison, we used the precomputed nuclear-solidity IQR feature provided with the original MorphLink analysis without additional mean-centering. The spatial spline basis dimension was 10. Genes were ranked in descending order of their MorphLink Curve-based Pattern Similarity Index (CPSI) with this feature and their MorphoANOVA location-specific H&E effect statistic $F_{32}$, respectively. Both rankings included all 38 genes. Concordance between the two rankings was evaluated using Spearman's rank correlation. For spatial interpretation, we examined *B2M*, which had the highest $F_{32}$, and decomposed its fitted total H&E effect into slide-wide and location-specific H&E components.

### References


[1] 10x Genomics. Visium FFPE human ovarian cancer web summary. `https://cf.10xgenomics.com/samples/spatial-exp/1.3.0/Visium_FFPE_Human_Ovarian_Cancer/Visium_FFPE_Human_Ovarian_`

Cancer_web_summary.html, 2024. Accessed: 2024-07-04.

[2] Jing Huang, Chenyang Yuan, Jiahui Jiang, Jianfeng Chen, Sunil S Badve, Yesim Gokmen-Polar, Rossana L Segura, Xinmiao Yan, Alexander Lazar, Jianjun Gao, et al. Bridging cell morphological behaviors and molecular dynamics in multi-modal spatial omics with MorphLink. *Nature Communications*, 16(1):5878, 2025.

[3] Tianzhi Wu, Erqiang Hu, Shuangbin Xu, Meijun Chen, Pingfan Guo, Zehan Dai, Tingze Feng, Lang Zhou, Wenli Tang, Li Zhan, Xiaocong Fu, Shanshan Liu, Xiaochen Bo, and Guangchuang Yu. clusterProfiler 4.0: A universal enrichment tool for interpreting omics data. *The Innovation*, 2(3):100141, 2021.

[4] Xi Jiang, Shidan Wang, Lei Guo, Bencong Zhu, Zhuoyu Wen, Liwei Jia, Lin Xu, Guanghua Xiao, and Qiwei Li. iIMPACT: integrating image and molecular profiles for spatial transcriptomics analysis. *Genome Biology*, 25:147, 2024.
